# Top-Down Feedback Controls Spatial Summation and Response Gain in Primate Visual Cortex

**DOI:** 10.1101/094680

**Authors:** Lauri Nurminen, Sam Merlin, Maryam Bijanzadeh, Frederick Federer, Alessandra Angelucci

**Author notes:** These authors contributed equally to this work (Lauri Nurminen, Sam Merlin). Current address: Department of Neurological Surgery, UCSF, CA 94143, USA. Current Address: Medical Science, School of Science and Health, Western Sydney University, Campbelltown, NSW 2560, Australia.

## Abstract

In the cerebral cortex, sensory information travels along feedforward connections through a hierarchy of areas, which, in turn, send a denser network of feedback connections to lower-order areas. Feedback has been implicated in attention, expectation, and sensory context, but the cellular mechanisms underlying these diverse feedback functions are unknown. Using specific optogenetic inactivation of feedback connections in the primate visual cortex, we have identified the cellular mechanisms of feedback-mediated modulations of early sensory processing. Specifically, we found that feedback modulates receptive field size, surround suppression and response gain, similar to the modulatory effects of visual spatial attention. A recurrent network model captured these effects. These feedback-mediated modulations allow higher-order cortical areas to dynamically regulate spatial resolution, sensitivity to image features, and efficiency of coding natural images in lower-order cortical areas.

---

In addition to well studied bottom-up feedforward inputs, the visual cortex receives a much denser network of feedback inputs from higher-order cortical areas^1^ whose role remains hypothetical. Feedback has been implicated in several forms of top-down influences, such as attention^2,3^, expectation^4^ and sensory context^5,6^, which affect sensory processing in diverse ways. For example, visual spatial attention, one of the most studied instances of top-down influences, has been shown to modulate neuronal response gain^2,7^, surround suppression^8^ and receptive field (RF) size^9^. In this study we have asked whether feedback connections can mediate such diverse effects.

To determine the cellular mechanisms underlying the influence of cortical feedback on sensory processing, we asked whether inactivating feedback from the secondary visual area (V2) alters RF size, surround suppression and response gain in the primary visual cortex (V1). Surround suppression is the property of V1 neurons to reduce their response to stimuli inside their RF when presented with large stimuli extending into the RF surround^10–18^. This is a fundamental computation throughout the visual cortex, thought to increase the neurons coding efficiency^19–22^, to contribute to segmentation of objects boundaries^21^, and to be generated by feedback connections^5,6^. However, the role of feedback in surround suppression and response gain remains controversial. Inactivation of higher-order cortices using pharmacology, cooling or optogenetics has produced weak reduction in surround suppression in some studies^23–25^, but only reduction in response gain in other studies^26–29^. One problem with these previous studies is that these inactivation methods suppressed activity in an entire cortical area; thus, the observed effects could have resulted from indirect pathways through the thalamus or other cortical areas. Moreover, these approaches did not allow fine control of inactivation levels, thus precluding potentially more physiologically relevant manipulations. To overcome the technical limitations of previous studies, we have used selective optogenetic inactivation of V2-to-V1 feedback terminals, rather than direct inactivation of the entire V2, while measuring spatial summation and surround suppression in V1 neurons using linear electrode arrays.

## RESULTS

### Specific Optogenetic Inactivation of Feedback Connections

To express the outward proton pump Archaerhodopsin-T (ArchT)^30^ in the axon terminals of V2 feedback neurons, we injected into V2 of marmoset monkeys a mixture of Cre-expressing and Cre-dependent adeno-associated virus (AAV9) carrying the genes for ArchT and green fluorescent protein (**Fig. 1a,c**; see Methods). This viral vector combination was used because in pilot studies we found it produces selective anterograde infection of neurons at the injected V2 site, and virtually no retrograde infection of neurons in V1 (**Fig. 1d**). Intrinsic signal optical imaging was used to identify the V1/V2 border (**Fig. 1a-b**) and target injections to V2 (**Fig. 1c-d**) (see Methods). Linear array recordings were, subsequently, targeted to GFP/ArchT-expressing V1 regions (**Fig. 1c,e**). Trial interleaved and balanced surface laser stimulation of increasing intensity was applied to ArchT-expressing axon terminals of V2 feedback neurons at the V1 recording site (**Fig. 1c**; see Methods). This viral injection protocol produces ArchT-GFP expression in V2 neurons at the injected site, including neurons sending feedback projections to V1 but also other V2 neurons projecting within V2 itself or to other brain regions. However, directing the laser to V1, while shielding V2 from light, allowed us to selectively inactivate V2 feedback terminals, leaving neurons within V2 unperturbed (**Fig. 1c**). Surface photostimulation most likely led to predominant inactivation of V2 feedback terminals in the superficial layers, and less so of feedback terminals in the deep layers of V1.

**Figure 1.**
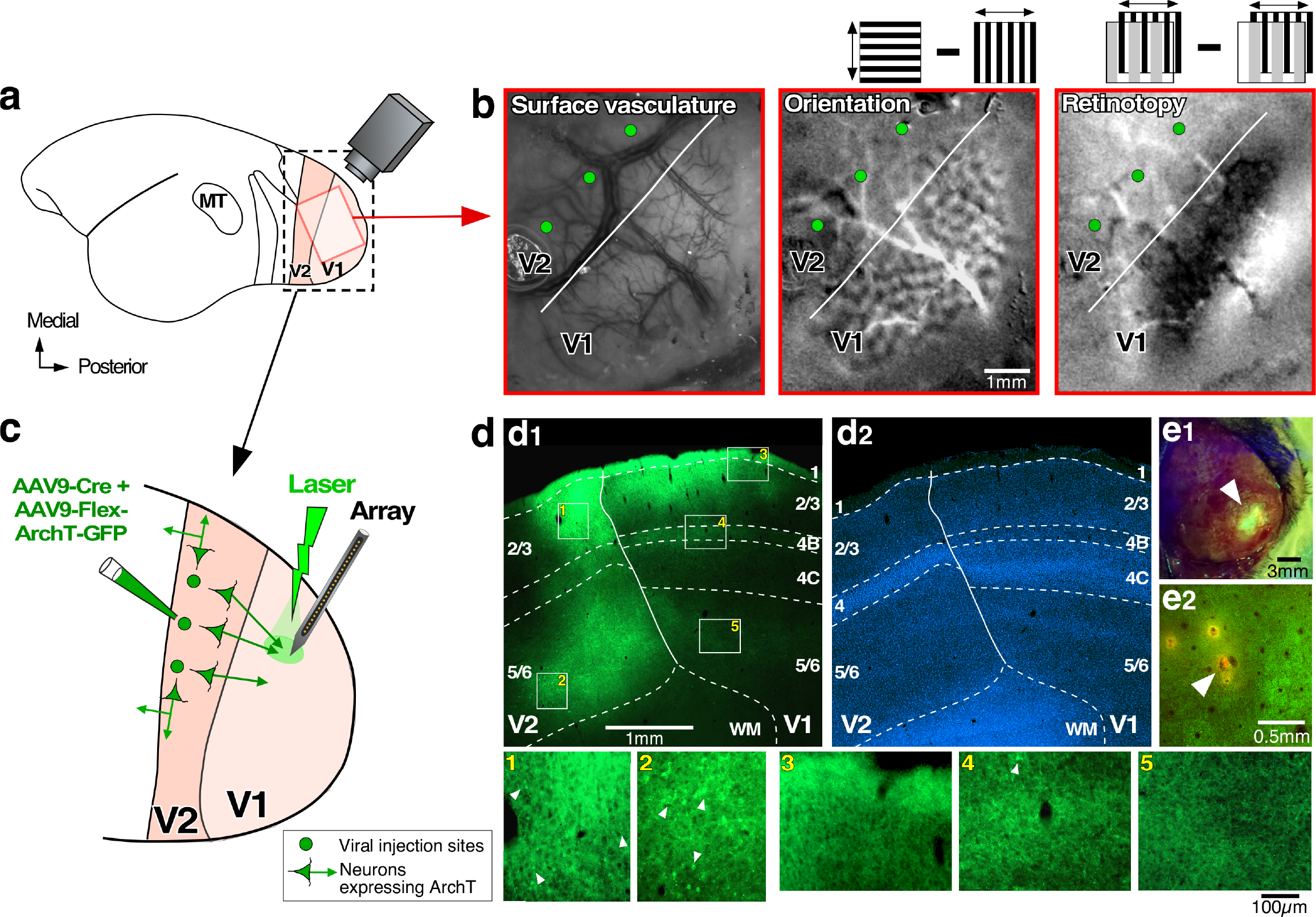
Optogenetic inactivation of V2 feedback terminals: experimental design and ArchT-GFP expression in V2 feedback terminals. **(a)** Schematics of lateral view of the marmoset brain. *Red box*: approximate location of the optically-imaged region in (b). *Black box*: V1 and V2 region shown enlarged in (c). **(b)** Optical imaging identifies V1/V2 border (*white line*). <u>Left panel:</u> cortical surface vasculature imaged under green light, used as reference to position pipettes for viral injections (*green dots*). <u>Middle panel:</u> differential orientation map generated by subtracting responses to gratings of 0°and 90° orientation (as shown in *inset*). V2 can be identified by larger orientation domains compared to V1. <u>Right panel</u>: retinotopic map generated by subtracting responses to 90° oriented gratings occupying complementary and adjacent strips (1°in width) of visual space (as shown in *inset;* see Methods). The V1/V2 border can be identified by the presence of stripes in V1, running approximately parallel to the V1/V2 border, which are absent in V2 (as the grating parameters were optimized for V1, but not V2, cell; see Methods). **(c)** Schematics of the inactivation paradigm: multiple viral injections were targeted to V2, array recordings and laser photostimulation to V1. **(d)** ArchT-GFP expression in V1 and V2. **(d_1_)**: Sagittal section through V1 and V2, viewed under GFP fluorescence, showing two viral injection sites confined to V2, and resulting expression of ArchT-GFP in the axon terminals of V2 feedback neurons within V1 layers 1-3, 4B and 5/6 (typical feedback laminar termination pattern^32,33^). This tissue section was located near the lateralmost aspect of the hemisphere, therefore the infragranular layers are elongated due to the lateral folding-over of the cortical sheet. *Solid contour*: V1/V2 border. *Dashed contours*: laminar borders delineated on the same section counterstained with DAPI **(d_2_)**. <u>BOTTOM (panels 1-5)</u>: higher magnification of label inside the *white boxes numbered 1-5* in (d_1_). Panels 1-2 show multiple clusters of labeled somata (e.g. *arrowheads*) at the V2 injection sites; instead, there is only one labeled soma (*arrowhead*) in panel 4, and none in panels 3,5. **(e_1_)** GFP excitation (*arrowhead*) through the intact thinned skull, approximately two months after viral injection. **(e_2_)** Tangential section through V1 showing the location of a DiI-coated electrode penetration (*arrowhead*) amid ArchT-GFP-expressing feedback axon terminals (*green fluorescence*).

### Feedback Affects Receptive Field Size

Electrophysiological recordings were performed in parafoveal V1 of anesthetized and paralyzed marmosets using 24-contact linear electrode arrays inserted orthogonal to the cortical surface, as verified by the vertical alignment of RFs and similarity of orientation preference across the array (see Methods, Supplementary Results and **Supplementary Fig. 1**). After initial characterization of RF properties at each contact through the V1 column, we measured spatial summation curves, using drifting grating patches of increasing diameter centered on the column’s aggregate RF. Typical V1 cells increase their response with stimulus diameter up to a peak (the RF size), and are suppressed for larger stimulus sizes activating also the RF surround (**Fig. 2a**). We extracted RF and surround diameter directly from the empirically measured size tuning curves as well as from phenomenological model fits to the size tuning data (see Methods), as described below.

**Figure 2.**
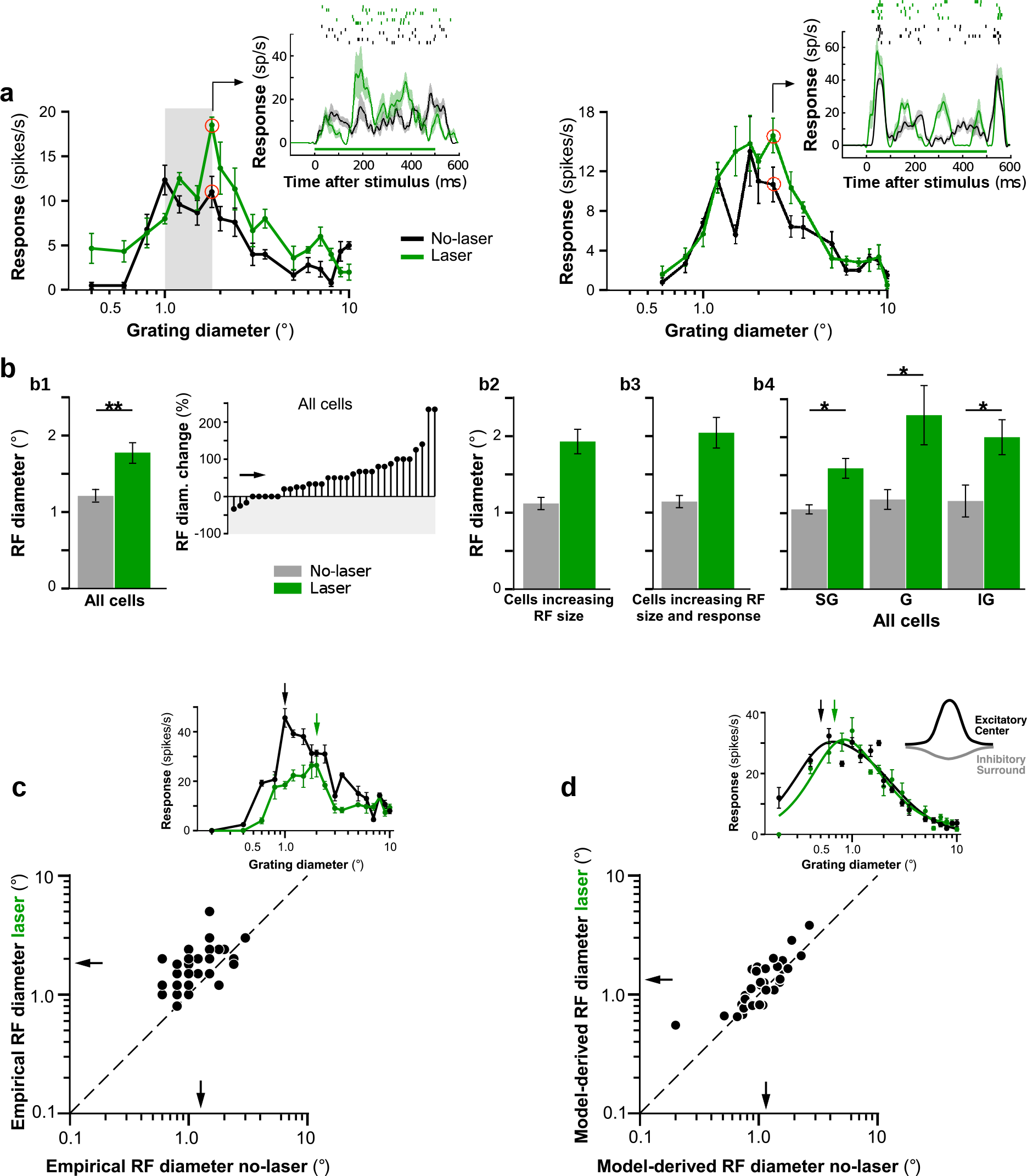
V2 feedback controls RF size. **(a)** Spatial summation curves for two example V1 cells recorded with (*green*) and without (*black*) laser stimulation. *Gray area in left panel*: proximal surround. *Insets*: PSTHs (BOTTOM; due to the smoothing filter used, response onset starts at time zero) and raster plots (TOP) measured at the stimulus diameters indicated by the *red circles* in the respective size-tuning curves. *Green horizontal line*: laser-on time. Two additional example cells are shown in the insets of panels (c) and (d). **(b)** Mean RF size (diameter at peak response of empirically-measured spatial summation curve) with and without laser stimulation for: **(b_1_)** All cells (LEFT; n=33); RIGHT: Cell-by-cell percent change in RF size across the entire cell population. Downward and upward stem: decreased and increased RF size, respectively. *Arrow*: mean. **(b_2_)** Only cells showing increased RF size with laser stimulation (n=25; mean RF diameter±s.e.m. no laser vs. laser: 1.12±0.08° vs. 1.93±0.08°). **(b_3_)** Only cells showing both increased RF size and peak response with laser stimulation (n=12; 1.14±0.08° vs. 2.04±0.20°). **(b_4_)** Mean RF size for all cells (as in b_1_), but grouped according to layer. *SG*: supragranular; *G*: granular; *IG*: infragranular. **(c-d)** Scatterplots of RF diameter with and without laser stimulation for RF diameter derived directly from the empirically measured summation curves (c), or from the model-fitted curves to the summation data (d), as indicated in the *insets* above each scatterplot. Insets in (c) and (d) show the size tuning curve of two additional example cells. The summation data for the cell in (d) are fitted with the ROG model. *Arrows* in insets in (c) and (d) indicate the RF diameters. *Arrows* in scatterplots: means. *Dashed line in (c-d)*: unity line.

We present spatial summation measurements from 67 visually responsive and stimulus modulated, spike-sorted single units from 3 animals. Approximately 61% (41/67) of single units were significantly modulated by the laser (see Methods, for neuronal sample selection). As laser-induced heat can alter cortical spiking activity^31^, we selected a safe range of laser intensities (9-43 mW/mm^2^), based on results from control experiments in cortex not expressing ArchT (see Supplementary Results and **Supplementary Figs. 2–3**).

When feedback was inactivated, the majority (76%) of laser-modulated units showed a shift of the spatial summation peak towards larger stimuli, i.e. an increase in RF size (**Fig. 2**); in the remainder of the cells RF size was unchanged (15%) or decreased (9%). Moreover, in 46% of cells RF size increase was accompanied by an increase in peak response amplitude (**Fig. 2a**), while in other cells peak response was decreased (e.g. **Fig. 2c** inset) or unchanged (e.g. **Fig. 2d** inset). This analysis was based on selecting, for each cell, the laser stimulation intensity producing the largest change in RF size, but within the range of intensities selected on the basis of control experiments (see above and Methods) (mean irradiance across the population ± sem was 28.7±1.95mW/mm^2^). Across the entire neuronal population (n=33 cells), mean RF diameter, defined as the stimulus diameter at the peak of the empirically measured summation curve (**Fig. 2c** inset), was significantly smaller with intact feedback, compared to when feedback was inactivated (mean±s.e.m: 1.27±0.10° vs. 1.83±0.14°, T-test p<0.01; Mann-Whitney U-test p<0.001; see Methods), with a mean increase of 56.2±10.7% (T-tes for mean increase >0%, p<0.001; Mann-Whitney U-test, p<0.001; **Fig. 2b_1_,c**). **Figure 2b2-b3** illustrates the magnitude of the mean RF size change caused by feedback inactivation, when considering only cells that showed increases in RF size (**Fig. 2b_2_**) or cells that showed increases in both RF size and peak response magnitude (**Fig. 2b_3_**).

We also examined how these changes in RF size vary with V1 layer. This was motivated by knowledge that V2 feedback connections show layer specific termination patterns in V1, targeting supragranular and infragranular layers, but avoiding the granular layer^32,33^. We found that feedback inactivation increased mean RF diameter in all layers (**Fig. 2b_4_**) (mean±s.e.m no-laser vs. laser: supragranular layers 1.23±0.11° vs. 1.53±0.10°; granular layer 1.31±0.17° vs. 2.26±0.35° infragranular layers 1.29±0.25° vs. 1.88±0.26°; T-test p<0.05 for all layers; Mann-Whitney U-test, p < 0.05 for all layers). This suggests that, at least in the granular layer, which does not receive direct feedback terminations, changes in RF size are relayed via other layers.

Since RF size derived from the empirically measured curves can be subject to noise, we also compared the RF size with and without laser extracted from phenomenological model fits to the summation data, as these can provide more robust measures of RF size. To this purpose, we fitted to the summation data two different models, namely a ratio or difference of integrals of two Gaussians (ROG or DOG model, respectively; see Methods), as these models have previously been shown to provide a good description of spatial summation curves in macaque V1^14,15^. In these models, a center excitatory Gaussian, corresponding to the RF center, overlaps a spatially broader inhibitory Gaussian, representing the suppressive surround (see inset in **Fig. 2d**); the major difference between the two models is that the surround inhibits the center through division in the ROG model, but through subtraction in the DOG model (see Methods). The ROG model provided a better fit for most (79%), but not all, of the cells (see below). Therefore, we fitted both models to the spatial summation data with and without laser stimulation, and for each cell we extracted RF size from the model that provided the best fit to that cell’s data. From the fitted curve, RF size was defined as the stimulus diameter at 95% of peak response (as in^14^) (**Fig. 2d** inset). Importantly, we still found feedback inactivation to significantly increase RF size when the latter was estimated from the models fits (**Fig. 2d**; mean diameter±s.e.m. no laser vs. laser: 1.15±0.09° vs. 1.34±0.12°, T-test p<0.01; Wilcoxon signed rank test p<0.05).

Additional analysis further demonstrated that increased RF size after feedback inactivation could not arise by chance, due to noise in the data (see Supplementary Results and **Supplementary Fig. 4**).

As feedback connections have been implicated in surround suppression, we asked whether inactivating feedback also affects the size of the RF surround. We found that whether derived from the empirical summation data or model fits to these data, the size of the surround field (see Methods for definition) was not affected by feedback inactivation either across the population (T-test p=0.33), or in individual layers (T-test p>0.27 for all layers) (see Supplementary Results). Because feedback connections from areas V3 and MT, which are spatially more extensive than feedback from V2^34^, were unperturbed in our study, a plausible explanation for this result is that feedback connections from these areas still provide large surround fields to V1 cells when V2 feedback is inactivated.

### V2 Feedback Affects Response Amplitude in the RF and Proximal Surround

Stimuli extending into the proximal surround (i.e. the surround region closest to the RF, here defined as the stimulus diameter at the peak of the laser-on size tuning curve, e.g. **Fig. 2a** left panel), evoked larger neuronal responses (mean±s.e.m. no-laser vs. laser: 36.4±12.3 vs. 43.5±17.2 spikes/s; mean increase 29.2±7.14%, T-test p<0.001; **Fig. 3a_1_**), and, therefore, less surround suppression (or even facilitation) with feedback inactivated when compared with intact feedback. Laser stimulation reduced the suppression index (SI; see Methods) for stimuli covering the RF and proximal surround, measured relative to the peak response in the no-laser condition (SI no-laser vs. laser: 0.21±0.03 vs. 0.006±0.0567, T-test p<0.01; **Fig. 3a_2_**). In contrast, the responses (no-laser vs. laser: 20.9±8.71 vs. 19.79 ±7.69 spikes/s; mean spike-rate decrease 7.10±13.4%, T-test p=0.92) and SI (no-laser vs. laser: 0.58±0.05 vs. 0.58±0.05; T-test p=0.945; **Fig. 3a_3_**) evoked by stimuli extending into the more distal surround were unchanged by feedback inactivation. V2 feedback inactivation is, indeed, expected to affect most strongly the suppression arising from the proximal surround, and to not abolish the most distal surround suppression. This is because feedback connections from V2 do not extend into the most distal surround regions of V1 neurons, unlike feedback connections from areas V3 and MT^34^, which were unperturbed in this study. Thus, the fact that the strength of surround suppression was mostly unaffected at the largest stimulus diameters is consistent with the known anatomical extent of feedback connections to V1 arising from different extrastriate areas^34^.

**Figure 3.**
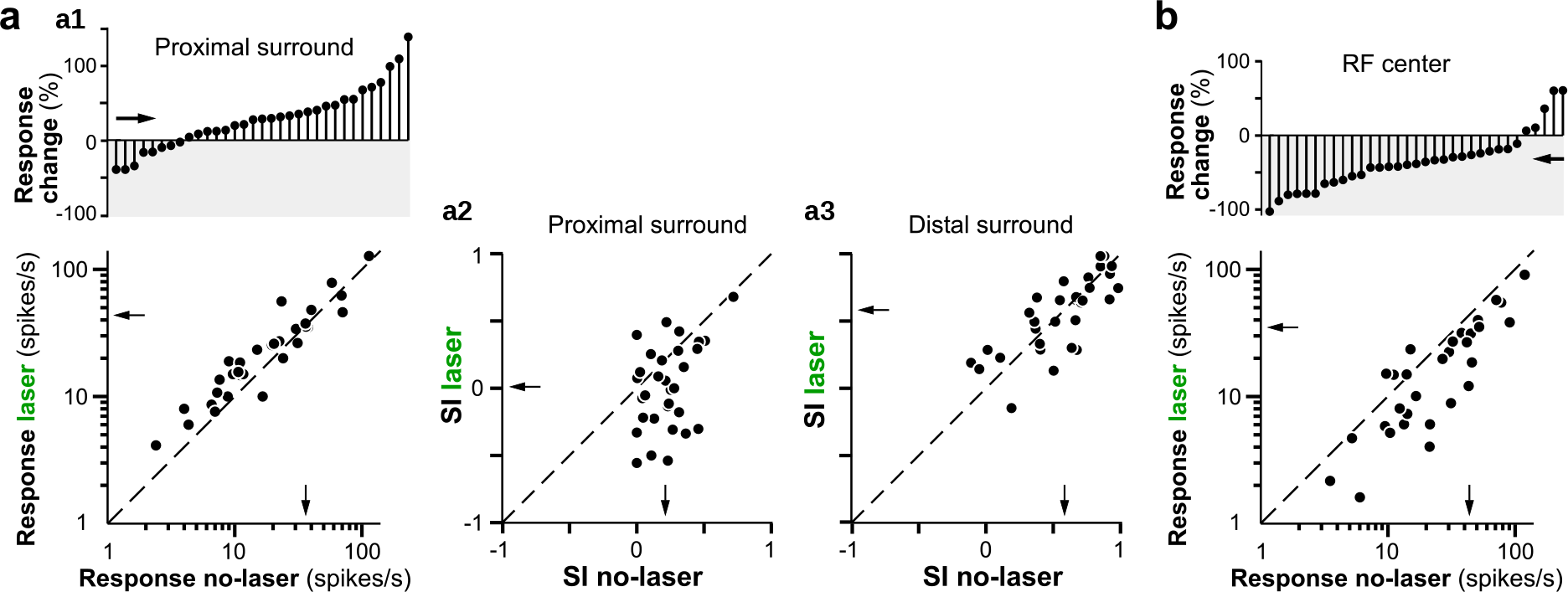
V2 feedback controls response amplitude in the RF and proximal surround. **(a)** Changes in proximal surround-suppression with V2 feedback inactivated. **(a_1_)** BOTTOM: response with and without laser for stimuli involving the RF and proximal surround. TOP: Cell-by-cell percent response change caused by laser stimulation, for stimuli extending into the proximal surround. Downward and upward stem: decreased and increased response, respectively. **(a_2_)** Suppression Index (SI; see Methods) with and without laser for stimuli extending into the proximal surround. SI=1 indicates maximal suppression, SI=0 indicates no suppression, and negative SI values indicate facilitation. **(a_3_)** Same as (a_2_) but for stimuli extending into the distal surround (largest stimulus used). **(b)** BOTTOM: response with and without laser for stimuli matched in size to the RF diameter (i.e. the stimulus diameter at the peak of the empirically-measured, spatial-summation curve in the no-laser condition). TOP: Cell-by-cell percent response change caused by laser stimulation for stimuli matched to the RF diameter. *Arrows*: means. One data point with very high firing rate in the scatterplots of panels (a) and (b) was removed for visualization purpose, but it was included in the analysis. *Dashed line in (a-b)*: unity line.

For some (e.g. **Fig. 2a** left panel), but not all (e.g. **Fig. 2a** right panel) neurons, inactivating feedback also changed the neuron’s response to small stimuli, the size of the neuron’s RF or smaller. We quantified these effects across the neuronal population. Consistent with previous studies of V2 inactivation^27,29^, we found that across the population of cells, stimuli matched in size to the neurons’ RF diameter (i.e. the stimulus diameter at the spatial summation peak in the no-laser condition) on average evoked lower responses in the laser condition (35.1±15.3 spikes/s) compared to the no-laser condition (43.8±14.1; mean reduction 32.0±6.03%, T-test p<10^−5^; **Fig. 3b**). Therefore, feedback inactivation reduced the gain of V1 neuron responses to stimuli inside the RF. However, although in some cells feedback inactivation increased neural responses to the smallest stimuli that evoked no response in the no-laser condition (e.g. **Fig. 2a** left panel), this increase was not significant across the population (average spike-rate difference between laser and no-laser conditions 1.28 ± 0.67 spikes/s, T-test p=0.39). We also found a moderate, but statistically insignificant, relationship between response reduction to stimuli matched to the RF diameter and change in RF diameter when feedback was inactivated (r= −0.31, p=0.11, Pearson’s correlation), as well as between change in RF diameter and release from suppression in the proximal surround (r=0.32, p=0.08).

Prolonged light pulses directed on ArchT-expressing axon terminals have been shown to facilitate synaptic transmission, while ArchT is consistently suppressive for pulse widths of ≤ 200ms^35^. Thus, we also performed the analysis described above focusing only on the first 200ms of the response. The results of the original and shorter time-scale analyses were qualitatively and quantitatively similar (see Supplementary Results), thus indicating that the observed results were caused by feedback inactivation.

### V2 Feedback Affects Response Gain

The analysis above revealed that in addition to increasing RF size, feedback inactivation also affected neuronal response gain. For most cells, responses to stimuli in the RF were reduced. However, responses to stimuli extending into the surround were increased in some cells (**Fig. 2a**), but decreased in other cells (**Fig. 2c** inset). We asked whether different levels of laser intensity had different impact on V1 neurons’ response gain.

**Figure 4a-b** shows two example cells in which RF size progressively increased and response amplitude progressively decreased with increasing laser intensity. However, the cell in **Figure 4b** showed greater and overall response reduction, while for the cell in **Figure 4a** response reduction was more pronounced at smaller stimulus diameters. Across the population of cells (n=33) we found that 36% of neurons showed response reduction across the entire spatial summation curve, and these were the neurons in the population that showed strongest surround suppression in the no-laser condition (SI: 0.78±0.03.1% vs. 0.49±0.07%, T-test p<0.05).

We quantified how RF diameter and mean response amplitude varied with laser intensity. This analysis is based on a population of 14 cells for which at least two laser intensities (within the range selected on the basis of the control experiments described in **Supplementary Figs. 2–3**) induced significance changes in the spatial summation curve (ANOVA p<0.05); for each of these cells the analysis was performed at the lowest (range: 3-31mW/mm^2^) and highest (range: 18-43mW/mm^2^) intensity.

**Figure 4.**
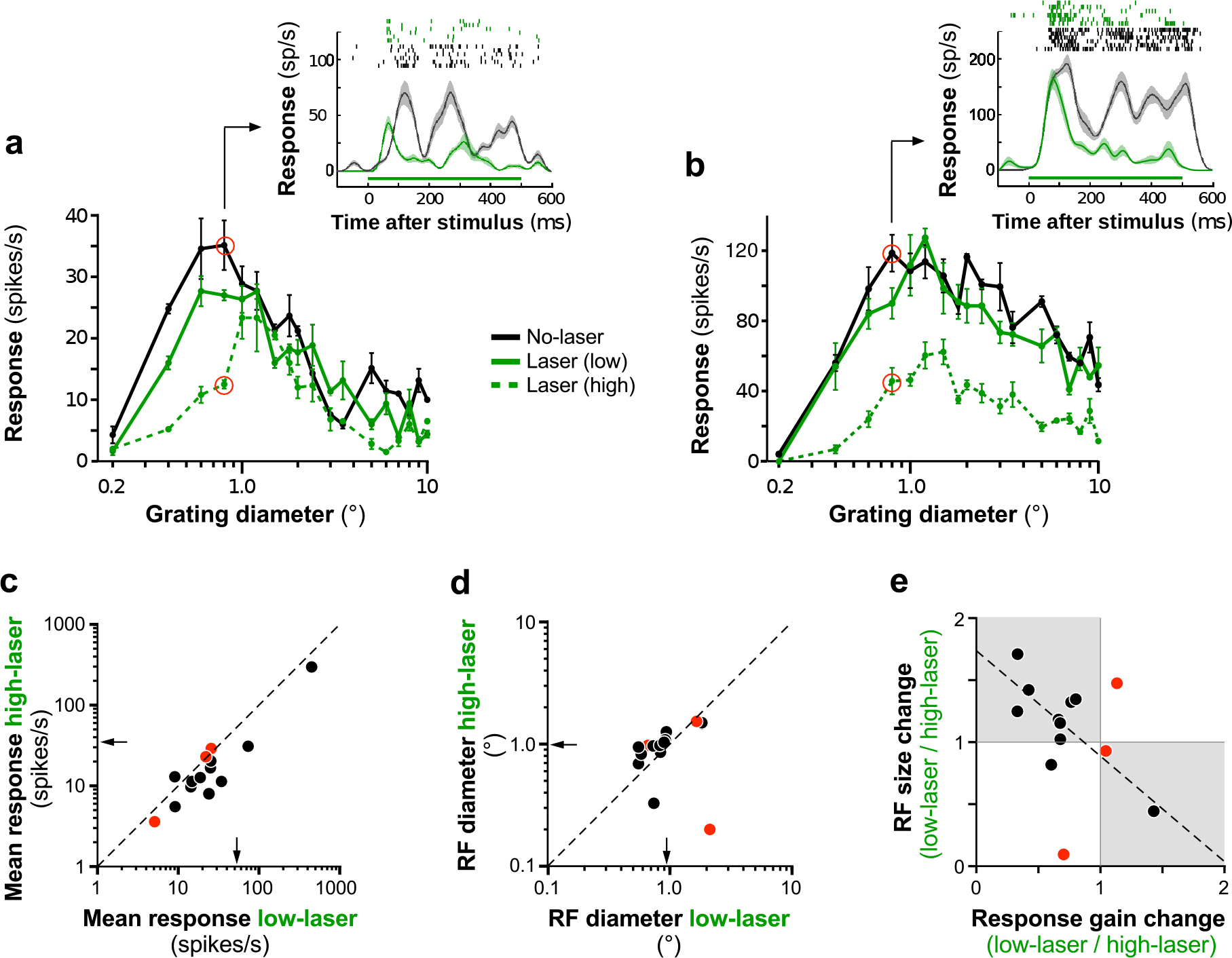
Feedback controls the gain of V1 responses. **(a-b)** Spatial-summation curves for two example V1 cells measured without laser (*black*) and with laser stimulation at two different intensities (*solid green*: 9 mW/mm^2^; *dashed green* 43 mW/mm^2^). Other conventions are as in **Fig.2a**. **(c)** Response gain (here defined as mean response over the entire spatial summation curve) for each cell at low and high laser intensity (n=14). **(d)** RF diameter for each cell at low and high laser intensity. *Black and red dots* in (c-e) indicate cells showing significant (T-test, p<0.05) and non-significant change in response gain, respectively. *Arrows* in (c-d): means. *Dashed line in (c-d)*: unity line. **(e)** RF size change (ratio of diameter at low to high laser) vs. response gain change (ratio of gain at low to high laser). *Shaded area* indicates cells for which RF size increased and response gain decreased with increasing laser intensity (and vice versa). *Dashed line*: regression line.

Compared to lower laser intensity, at higher laser intensity 11/14 cells showed a significant reduction in mean response amplitude (T-test p<0.05; **Fig. 4c**) and 10/14 cells showed increased RF diameter (**Fig. 4d**). Furthermore, most cells (10/14) showed both, reduced response gain and increased RF size with increasing laser intensity (**Fig. 4e**). For the cells that showed a statistically significant gain change at higher laser intensity (n=11; black dots in **Fig. 4e**), there was a significant negative correlation between response gain change and RF size change (r= −0. 77, Pearson’s correlation, p<0.01).

### Mechanisms Underlying the Effects of Feedback Inactivation

We used both phenomenological and network models, to gain insights into the mechanisms that underlie the effects of feedback inactivation on RF size and response gain.

Because the RF and surround interactions have been typically modeled as two overlapping but distinct Gaussian mechanisms interacting either divisively (ROG model) or subtractively (DOG model)^14,15^, we fitted both a ROG and a DOG model to the spatial summation data presented in **Figure 2** in the laser and no-laser conditions, and compared how well each model fitted the data (see Methods). We found that for the majority of the cells (79%) the ROG model provided a better fit to the data (mean R^2^±s.e.m. for cells that were best fit by the ROG model 0.67±0.04 vs. 0.37±0.10 for the DOG model fits to the same cells). For the reminder of cells (21%), both models provided similar good fits to the data. This result is consistent with the idea that the surround affects neural responses with and without feedback via divisive normalization mechanisms^14^.

We next determined which model parameters were mostly affected by feedback inactivation. To achieve this goal, we selected for each cell the model that provided the best fit to its size tuning measurements in the no-laser condition, and then allowed one parameter at a time to vary with feedback inactivation, while the remainder of the model parameters were held fixed to their original values. As none of the single parameter models could account for the full range of the effects seen in the inactivation data (see Supplementary Results and **Supplementary Fig. 5a**), we next allowed two parameters at a time to vary with feedback inactivation, while holding the rest constant, and performed this analysis for all possible combinations of parameter pairs. The model in which both the spatial extent and gain of the center excitatory mechanism were allowed to vary best accounted for the inactivation results of 30% of cells in the population, followed by a model in which the spatial extent of both the excitatory center and inhibitory surround mechanisms were varied, which, instead, provided best fits for 21% of the cells (see Supplementary Results and **Supplementary Fig. 5b-c**). However, none of the two-parameter models provided best fit for the majority of the cells. Moreover, when comparing the different models based on the coefficient of determination (R^2^) distributions, rather than fraction of cells best fit by each model, we found that the different models performed similarly (see Supplementary Results and **Supplementary Fig. 5d**).

To gain better insights into the circuit mechanisms underlying changes in RF size and response gain induced by feedback inactivation, we used neural network modeling. As the effects of feedback inactivation on spatial summation are reminiscent of the effects of changing stimulus contrast^14,36^, we used a previously published recurrent network model of V1, which well described the spatial summation properties of V1 cells and their contrast dependence. In particular, we asked whether similar circuit mechanisms underlying changes in RF size and gain with stimulus contrast could also account for the effects of feedback inactivation.

We used the 1D recurrent network model of Schwabe et al.^37^, which accounts for surround suppression in V1 using intra-V1 horizontal and local recurrent connections, feedback connections from a single extrastriate area, and a single population of inhibitory (I) neurons (**Fig. 5a**; see Methods). In this model, I neurons have higher threshold and gain than excitatory (E) neurons (**Fig. 5b**) and, consistent with recent findings^38^, are more strongly driven by horizontal connections than the E cells whose output they control. As a result, I cells generate suppression under sufficiently high levels of excitation, but are inactive for low levels of excitation. Therefore, the local network in the model becomes more dominated by inhibition with increasing excitatory drive. For weak excitatory inputs (e.g. small visual stimuli in the RF), I neurons are silent, but for strong inputs (e.g. large stimuli encompassing the RF and surround), they become active (**Fig. 5c** *dashed pink curve*) and suppress the E neurons’ response (**Fig. 5c** *black curve*). Therefore, the model I neurons behave similarly to somatostatin neurons in mouse visual cortex^38^, beginning to respond at larger stimulus sizes than E neurons, and increasing their response with increasing stimulus size, thus causing surround suppression.

**Figure 5.**
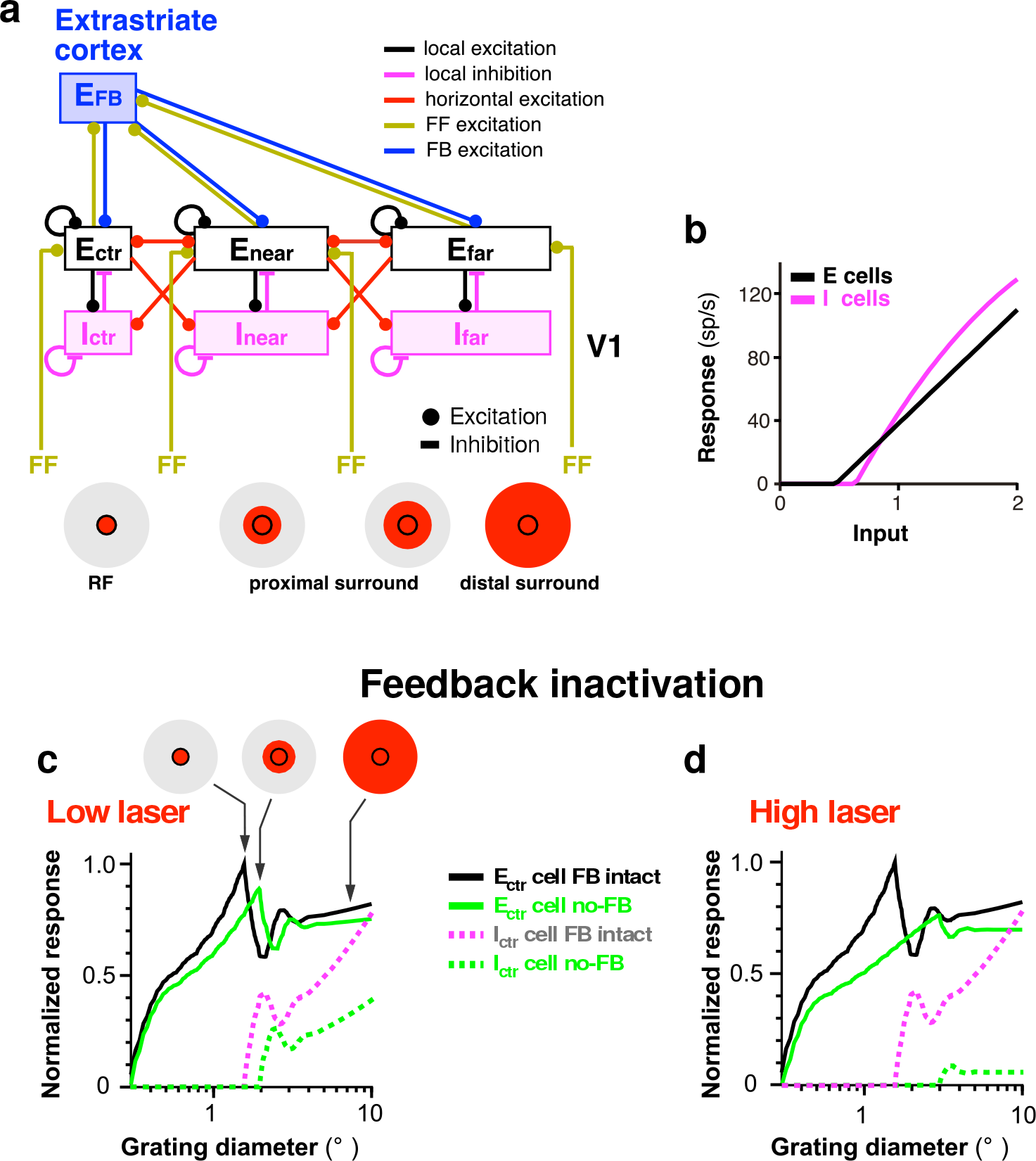
Effects on spatial summation of inactivating feedback connections in a recurrent network model of V1. **(a)** The recurrent network model architecture. Diagram of the connections in the model. Different connection types are color coded according to the legend. *Pink and black boxes*: population of layer 2/3 inhibitory (I) and excitatory (E) cells, respectively, labeled according to the position of their RFs relative to that of the cells in the center recorded V1 column; accordingly, E_ctr_/I_ctr_ are the cells in the center column, and E_near, far_/ I_near, far_ are the cells in the near and far surround, respectively. The near surround in the original model was defined as the surround region co-extensive with the spatial spread of V1 horizontal connections, and the far surround as the region beyond it, coextensive with the spread of feedback connections from the extrastriate area. The *proximal surround*, as defined in this study, encompasses the near surround and the more proximal region of the far surround, while the *distal surround* encompasses the more distal region of the far surround. *FF*: excitatory feedforward afferents from other V1 layers to layers 2/3; *E_FB_*: excitatory feedback connections from a single extrastriate area to V1. *Icons* at the bottom and in panel (c) represent RF and surround components, with red areas indicating regions that are activated by a stimulus of increasing diameter. **(b)** Firing rate of the local E and I cells in the model, plotted against the input current. **(c, d)** Size tuning curves of the model Ectr and Ictr neurons with intact feedback and with low (c) or high (d) levels of feedback inactivation. Low levels of feedback inactivation (obtained in the model by multiplying the feedback weights by 0.5) reduces neuronal responses to small stimuli inside the RF, increases RF size and increases responses in the proximal surround (similar to the results shown in **Figs. 2-3**. **(d)** High levels of feedback inactivation (feedback weights set to 0) further increases RF size and reduces response gain, similar to the results shown in **Fig. 4**.

We found that a similar mechanism as that underlying the increase in RF size at low stimulus contrast, in this model, also accounts for the increase in RF size when feedback is inactivated. Specifically, in the model, moderate reduction of feedback excitation to the V1 network, weakens the response of I neurons (**Fig. 5c** *dashed green curve*), allowing E neurons to summate excitatory signals over larger visual field regions (i.e. to increase their RF size; **Fig. 5c** *solid green curve*) until the I neurons’ threshold is reached leading to suppression of E neurons (**Fig. 5c** *green curves*). Further reducing feedback excitation, as achieved by increasing laser intensity, leads to both further increase in RF size and further decrease in response gain (**Fig. 5d** *solid green curve*). This is consistent with the behavior of most cells in **Fig. 4e** (data points in the shaded squares), for which we indeed found a significant negative correlation between RF size change and response gain change when laser intensity was increased. Therefore, a single mechanism in the network model can account for the main effects of feedback inactivation, i.e. increased RF size and response gain change.

The network model could not easily reproduce the overall strong reduction in response amplitude of the entire summation curve, as seen in 36% of cells, particularly at higher laser intensity (e.g. **Fig. 4b**), perhaps because it relies on a single inhibitory neuron type. Moreover, V1 receives feedback connections from multiple extrastriate areas, whose spatial extent increases with the area’s hierarchical distance from V1^34^. As the model incorporates feedback connections at a single spatial scale, it cannot optimally reproduce the differential effects on proximal vs. distal surround suppression of removing feedback from a single area, while leaving intact more extensive feedback from other areas. Specifically, far surround suppression in the model was weaker than in the data. Thus, future refinements of this model will have to incorporate feedback at multiple spatial scales and multiple inhibitory neuron types.

## DISCUSSION

Our study elucidates how feedback affects neural responses in the primate early visual cortex. Reducing V2 feedback activity increased RF size, decreased V1 cell’s responses to stimuli confined to their RF, and increased their responses to stimuli extending into the proximal surround, thus weakening surround suppression. The magnitude of these effects depended on the degree of feedback inactivation, so that stronger reduction of V2 feedback activity led to greater increase in RF size and progressive decrease in response amplitude. Therefore, our results indicate that feedback from V2 controls RF size, proximal surround suppression and response gain in V1.

Our study is the first to demonstrate that feedback is part of the network that regulates the RF size of V1 neurons. None of the previous studies reported systematic effects of inactivating extrastriate cortex on V1 cells’ RF size^23–29^. For most of these previous studies, this is because RF size was not measured after inactivation of higher cortical areas^23,24,26,27,29^. In two prior studies^25,28^, however, spatial summation measurements similar to those performed in our study were made before and after inactivation of higher visual cortex. It is unclear why no systematic effects of inactivating extrastriate cortex on RF size were observed in these two studies, but differences with our study that could have led to the different results include the specific cortical areas that were inactivated (macaque V2 and V3 ^25^, or cat postero-temporal visual cortex^28^, likely homologue of macaque inferotemporal cortex), inactivation methods (cooling of entire cortical area/s), and data analysis. Compared to previous studies, which silenced an entire cortical area, therefore also affecting activity in downstream cortical or subcortical areas, the strength of our approach is the selective and titrated manipulation of feedback neuron activity. This may have allowed us to reveal nuanced effects caused selectively by direct feedback to V1, which could have been missed with coarser cooling methods.

Consistent with our findings, most previous studies in anesthetized animals have reported that inactivating extrastriate cortex leads to reduced responses to stimuli inside the RF of V1 cells^23,24,27–29^. In contrast, cooling areas V2 and V3 simultaneously in awake primates produced variable effects on the magnitude of V1 RF responses, including increases and decreases^25^; this variability may have been caused by fixational eye movements, to which the small RFs of V1 neurons are particularly sensitive.

There has been lack of consensus over which circuits generate surround suppression in V1, in particular whether this is generated subcortically and relayed to V1 via geniculocortical connections, or intracortically by V1 horizontal connections and/or feedback connections from extrastriate cortex. Current experimental evidence suggests that all these connection types, in fact, contribute to surround suppression in V1^5^. On the one hand, suppression in V1 caused by large stimuli can occur as fast as visual responses to RF stimulation^39,40^, and first emerges in V1 geniculocortical input layer 4C^41^; moreover, this early suppression is untuned for stimulus orientation^39,41^. These findings suggest that the earliest untuned suppression in V1 is inherited from the lateral geniculate nucleus, where neurons also show untuned surround suppression^42,43^. On the other hand, two recent optogenetic studies in mouse have provided direct evidence for a contribution of intra-V1 horizontal connections to surround suppression in V1^38,44^.

A role for feedback in surround suppression was suggested on the basis of evidence that feedback, but not monosynaptic horizontal, connections encompass the full spatial extent of the RF and surround of V1 neurons^34^, and conduct signals 10 times faster than horizontal axons^45^. Thus, the slower conduction velocity and limited spatial extent of horizontal connections would seem inadequate to mediate fast suppression^46^ arising from the more distal regions of the surround of V1 neurons^6^. However, previous inactivation studies have provided contrasting results regarding the role of feedback in surround suppression. Some studies observed weak reduction in surround suppression after cooling primate area MT^23^ or V2 and V3 together^25^, or cat postero-temporal visual cortex^24^. Other studies, instead, found general reduction in response gain, but no change in surround suppression after pharmacologically silencing primate V2^27^, cooling cat postero-temporal visual cortex^28^ or optogenetically silencing mouse cingulate cortex^26^. In our study, feedback inactivation caused both reduced surround suppression and changes in response gain, with reduced response gain most often observed after stronger feedback inactivation. Therefore, our results support the involvement of feedback in both surround suppression and response gain. The discrepancy between studies on the effects of feedback inactivation on surround suppression could be attributed to several differences, including levels and spatial extent of feedback inactivation, the specific cortical area inactivated (two of these studies inactivated higher level cortical areas), and methods of quantifying surround suppression that did not take into account the spatial extent of the specific feedback system that was inactivated.

Inactivating V2 feedback reduced suppression predominantly in the proximal surround, and did not abolish distal surround suppression. This was predicted on the basis of the known visuotopic extent of V2 feedback connections. The latter are less extensive than feedback connections arising from areas V3 and MT, which, instead, encompass the full extent of the distal surround^34^. What may be surprising is that inactivating V2 feedback had no effects on distal surround suppression. Linear summation predicts a reduction (but not abolishment) of suppression caused by the largest stimuli when V2 feedback inactivation leads to reduced proximal surround suppression. Therefore, our finding suggests that feedback from different extra-striate areas affect V1 responses via a common non-linear mechanism.

To gain insights into the mechanisms underlying the impact of feedback on V1 neuron responses, we fitted the data with phenomenological models previously used to describe the effects of contrast on RF size^14,36^, as well as the effects of inactivating areas V2 and V3 on surround suppression in V1^47^. In these models, the RF and surround have Gaussian sensitivity profiles, with the RF described as an excitatory Gaussian and the surround as an inhibitory Gaussian, the two interacting either subtractively or divisively. Sceniak et al.^36^ found that at low stimulus contrast, RF size is larger and response gain is lower than at high contrast, and suggested this results from an increase in the spatial extent of the center Gaussian mechanism. Cavanaugh et al.^14^, instead, demonstrated that contrast-dependent changes in RF size and gain could be explained by changes in the gain of both the center and surround Gaussian mechanisms. Our modeling results differ from these previous reports, because although the effects of contrast on RF size and gain resemble some of the effects of feedback inactivation, particularly those we have observed at higher laser intensity, they nevertheless represent only a subset of the full range of feedback inactivation effects. Thus, models in which feedback inactivation modifies only the spatial extent of the center Gaussian^36^, or only the gain of both the center and surround Gaussians^14^ could capture the increase in RF size and gain reduction, but failed to capture the simultaneous response decrease to stimuli in the RF and response increase to stimuli extending into the proximal surround.

The modeling work of Nassi et al.^47^ showed that changes in the spatial extent of inhibition best accounted for changes in V1 spatial summation after simultaneous cooling of macaque areas V2 and V3. In agreement with this previous study, we found that such a model could capture the changes in neural responses for stimuli extending into the surround, i.e. the reduction in surround suppression found in both our and these authors’ study. However, in contrast to Nassi et al., we found that feedback inactivation also caused an increase in RF size, and this could not be accounted for by a model in which feedback only affects the spatial extent of surround inhibition. Instead, we found that a model involving changes in the spatial extent and gain of the excitatory mechanism best accounted for the range of feedback inactivation effects, suggesting that V2 feedback affects the spatial extent over which cells integrate excitation as well as the gain of excitation.

A simple network model in which spatial summation results from the interaction of feedforward, V1 horizontal and inter-areal feedback connections with local recurrent networks, provided further insights into the network mechanisms that may underlie these effects of feedback inactivation. In this model, changes in RF size and response amplitude after feedback inactivation were explained by a single mechanism, asymmetric inhibition, which leads to an altered balance of excitation and inhibition when excitatory feedback inputs to E and I neurons are reduced. While in our model asymmetric inhibition is implemented using high-threshold/gain somatostatin-like inhibitory neurons, in principle other models with asymmetric inhibition should be able to account for feedback inactivation effects on RF size and response amplitude. For example, in the model of Rubin et al.^48^, asymmetric inhibitory/excitatory responses are implemented using a mechanism based on a supralinear input/output function of cortical neurons (which causes the gain of the input/output function to increase with increasing postsynaptic activity) and an inhibition-stabilized network (in which strong recurrent excitation is stabilized by strong recurrent inhibition). As this model can well explain contrast-dependent changes in RF size and gain, it will be interesting to see if it can also account for the variety of gain changes induced by feedback inactivation.

Finally, it is important to point out that several forms of top-down influences in sensory processing have been shown to affect neuronal responses in the same way as we have shown here for feedback from V2. For example, spatial attention increases the response gain of neurons at attended locations^2,7^, modulates surround suppression^8,49^ and, at least in parafoveal V1, modulates RF size^9^. Our results suggest that these effects can all be mediated by top-down modulations of feedback to early visual areas.

Together, our findings suggest that the function of V2 feedback is to control the spatial resolution of visual signals and the perceptual sensitivity to image features, by modulating the RF size and response gain of V1 neurons, respectively, and to contribute to surround suppression in V1.

## METHODS

### Surgery and Viral Injections

All procedures conformed to the guidelines of the University of Utah Institutional Animal Care and Use Committee. Each of three adult marmoset monkeys (*Callithrix jacchus*) received 2-3 injections in dorsal area V2 of a 1:1 viral mixture of AAV9.CaMKII.Cre (3.7×10^13^ particles/ml) and AAV9.Flex.CAG.ArchT-GFP (9.8×10^12^ particles/ml; Penn Vector Core, University of Pennsylvania, PA). Injections were targeted and confined to V2 using as guidance the location of the V1/V2 border identified *in vivo* using intrinsic signal optical imaging. Surgical procedures were as previously described^50^. Briefly, animals were pre-anesthetized with ketamine (25-30mg/kg, i.m.) and xylazine (1mg/kg, i.m.), intubated, artificially ventilated with N_2_O and O_2_ (70:30), and the head was stereotaxically positioned. Anesthesia was maintained with isoflurane (1-2%), and end-tidal CO_2_, blood oxygenation level, electrocardiogram, and body temperature were monitored continuously. The scalp was opened and the skull was thinned using a dental drill over areas V1/V2, covered with agar and a coverslip, which was glued to the skull. On completion of surgery, isofluorane was turned off, anesthesia maintained with sufentanil citrate (8-13μg/kg/hr, i.v.), and paralysis was induced with repeated 30-60 min intravenous boluses of rocuronium bromide (0.6mg/kg/hr) to stabilize the eyes. The pupils were dilated with a topical short-acting mydriatic agent (tropicamide), the corneas protected with gas-permeable contact lenses, the eyes were refracted, and optical imaging was started. Once the V1/V2 border was functionally identified, the glass coverslip was removed, small craniotomies and durotomies were performed over V2, and the viral mixture slowly pressure-injected (240nl/site at 500μm and again at 1200μm depth, using glass pipettes of 40-50μm tip diameter, 15 minutes/240nl). The thinned skull was reinforced with dental cement, the skin sutured and the animal recovered.

### Optical Imaging

Acquisition of intrinsic signals was performed using the Imager 3001 (Optical Imaging Ltd, Israel) under red light illumination (630 nm). Imaging for orientation and retinotopy allows identification of the V1/V2 border (**Fig. 1a-b**). Orientation maps were obtained using full-field, high-contrast (100%), pseudorandomized achromatic drifting square-wave gratings of 8 orientations at 0.5-2.0 cycles/° spatial frequency and 2.85 cycles/s temporal frequency, moving back and forth, orthogonal to the grating orientation. Responses to same orientations were averaged across trials, baseline subtracted, and difference images obtained by subtracting the response to two orthogonal oriented pairs (e.g. **Fig. 1b** middle panel). Retinotopic maps were obtained by subtracting responses to monocularly presented oriented gratings occupying complementary adjacent strips of visual space, i.e. masked by 0.5-1° strips of gray repeating every 1-2°, with the masks reversing in position in alternate trials (**Fig. 1b** right panel)^51^. In each case, reference images of the surface vasculature were taken under 546 nm illumination (green light, **Fig. 1b** left panel), and later used as reference to position pipettes for viral vector injection.

### Electrophysiological Recordings and Visual Stimulation

Following 62-68 days after the viral vector injection, animals were anesthetized and paralyzed by continuous infusion of sufentanil citrate (6-13μg/kg/h) and vecuronium bromide (0.3mg/kg/h), respectively, and vital signs were continuously monitored, as described above. The pupils were dilated with topical atropine, protected with lenses and refracted. GFP-expressing V2 injection sites and V2 feedback axonal fields in V1 were identified with GFP goggles (**Fig. 1e_1_**), and small craniotomies were made over V1. Extra-cellular recordings were made in V1 with 24-channel linear multielectrode arrays (V-Probe, Plexon, Dallas, TX; 100μm contact spacing, 20μm contact diameter) coated with DiI (Molecular Probes, Eugene, OR) to assist with post-mortem reconstruction of the electrode penetrations (e.g. **Fig. 1e_2_**), and lowered normal to the cortical surface (using triangulation methods) to a 2-2.2 mm depth over 60-90min. A 128-channel system (Cerebus, Blackrock Microsystems, Salt Lake City, UT) was used for signal amplification and digitization (30 kHz). Continuous voltage traces were band-pass filtered (0.5-14.25 kHz), and spikes were detected as spatiotemporal waveforms using the double-threshold flood fill algorithm^52^ (thresholds 2 and 4 × noise S.D.). This procedure was adopted because the apical dendrites of pyramidal cells run parallel to the probe shank and may spread the same waveforms across multiple channels. A masked EM algorithm^53^ was used for clustering, and manual refinement of the clusters was performed with the Klustasuite^52^.

After manually locating the recorded RFs, their aggregate minimum response field was quantitatively determined using a sparse noise stimulus (500ms, 0.0625-0.25 deg^2^ square, luminance decrement, 5-15 trials; **Supplementary Fig. 1b**) and all subsequent stimuli were centered on this field. Orientation, eye, spatial and temporal frequency preferences for the cells in the recorded V1 column were determined using 1° diameter, 100% contrast drifting sinusoidal gratings monocularly presented on an unmodulated gray background of 45cd m^−2^ mean luminance. We then performed spatial summation measurements using circular patches of 100% contrast drifting sinusoidal gratings of increasing diameter centered over the columnar aggregate minimum response field. The patch diameter ranged from 0.2-0.6° to 10-18° (depending on animal) and different patch sizes were presented in random order within each block of trials. All size-tuning experiments were performed using gratings of spatial and temporal frequencies and orientation that strongly drove most cells in the column. It was not typically challenging to find spatial and temporal frequency values to which all cells in the column responded vigorously. When the penetration was perfectly vertical, orientation preference was also similar for all cells in the column. Slight deviations from vertical, however, even for RFs perfectly aligned in space, could cause orientation to shift slightly across the column, due to the narrow orientation tuning of many V1 cells^54^. In this case, the size tuning experiment was run using two different orientations. Importantly, although deviations from optimal stimulus parameters can increase the neurons’ summation area^55^, these deviations are not expected to cause differences between neuronal responses recorded with and without laser stimulation. To monitor eye movements, the RFs were remapped by hand approximately every 10 minutes, and stimuli were re-centered in the RF when necessary. Stimuli were presented for 500ms with 750ms inter-stimulus interval. Stimuli were programmed with Matlab (Mathworks, Natick, MA) and presented on a linearized CRT monitor (Sony GDM-C520, 600 × 800 pixels, 100Hz, 57cm viewing distance) and their timing was controlled with the ViSaGe system (Cambridge Research Systems, Cambridge, UK). Data analysis was performed using custom scripts written in Matlab and Python^56,57^.

### Laser Stimulation

A 532nm laser (Laserwave, Beijing, China) beam was coupled to a 400μm diameter (NA=0.15) optical fiber, then expanded and collimated to a 2.8 mm spot. Reported irradiances refer to the light power exiting the collimator divided by the area of the collimator. Because the beam was collimated, the illumination spot size depended very little on the distance of the fiber from the brain. Laser timing was controlled at submillisecond precision, using custom made programs running on real-time Linux. Light was shone on the surface of V1 through thinned skull in the regions of GFP expression, and V2 was shielded from light. Laser onset was simultaneous with stimulus onset and photostimulation continued throughout stimulus presentation (500ms). The animal’s eyes were shielded from the laser light.

### Neuronal Sample Selection

We analyzed 67 visually responsive (defined as max response at least 2SD>baseline) and stimulus modulated (one-way ANOVA, p<0.05) units. Approximately 61% (41/67) of these visually driven single-units were modulated by one or more laser stimulation intensities (twoway ANOVA, either laser or stimulus diameter × laser interaction, p<0.05, or at least two successive data points different in the same direction, p<0.05). We were not able to determine RF size for eight cells, thus these were excluded from further analysis. Therefore, a total of 33 cells were analyzed for the results reported in **Figs. 2–3**. **Fig. 2b2-b3** are based on smaller populations of cells within this larger population of 33 cells (as indicated in the figure legend), and the three populations were not mutually exclusive.

For the analysis of the data presented in **Fig. 2**, the laser stimulation intensity producing the largest change in RF size (but within the range of intensities selected on the basis of control experiments-see **Supplementary Figs. 2–3** and Supplementary Results) was determined for each unit separately, and the analysis was performed at this intensity. This was motivated by expectations that the light intensity required to produce inactivation effects differs among cells due to several factors, including variation in opsin expression across neurons, distance of the cells from the light source, and intrinsic differences in sensitivity to feedback perturbation. Importantly, however, even though we selected different light intensities for different cells, the direction of the effects was not biased by our analysis, as we selected for each cell the laser intensity causing the largest change in RF size, irrespective of whether this was an increase or decrease.

The analysis of the data presented in **Fig. 4** is based on a population of 14 cells for which at least two laser intensities (within the range selected on the basis of the control experiments described in **Supplementary Figs. 2–3**) induced significance changes in the spatial summation curve (ANOVA for either laser or stimulus diameter × laser interaction p<0.05). This is a subset (14/33) of the population analyzed in **Figs. 2–3**, because for the remainder of the population we either lacked two laser intensity levels, or only one laser intensity (within the range selected on the basis of control experiments) caused significant changes.

### Definition of RF and Surround Size

From the size tuning curves, measured as described above, for each cell we extracted as a measure of RF size the grating’s diameter eliciting maximum response. Surround size was defined as the smallest grating diameter for which the neuron’s response was reduced to within 5% of the response at the largest diameter. As these measures of RF and surround size can be subject to noise, to derive more robust measures, we also fitted the size tuning data with the ratio and difference of the integral of two Gaussian functions (ROG^14^ and DOG^15^ models, respectively; see below for model fits). From the fitted summation curves we extracted the cells’ RF size as the smallest stimulus diameter at which the cell response reached 95% of the peak response^14^.

### Statistical Model Fitting

ROG^14^ (eq. 1) and DOG^15^ (eq. 2) models were fitted to the size tuning data according to the following functions

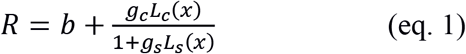

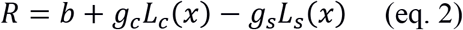

where

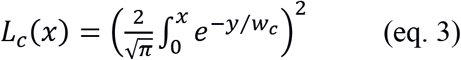

and

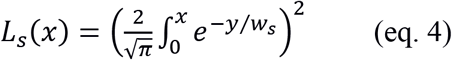

Here the variable × corresponds to the diameter of the stimulus, wc and ws are the spatial extents of the center excitatory and surround inhibitory Gaussian mechanisms, respectively (with the constraint that w_c_ < w_s_), L_c_ and L_s_ are the activities of the center and surround mechanisms, respectively, and gc and gs are the gains of the center and surround mechanisms, respectively. All parameters were constrained to positive values during optimization. Model parameters were optimized by minimizing the sum of squared errors between the model predictions and the data. Initial parameter search was done by performing two successive grid optimizations. The first grid was coarse, and the second grid was finely spaced and centered on the best fitting parameters determined with the first grid search. The best fitting parameters determined with the second grid were used as initial parameters for final optimization, which was done using the active-set algorithm in Matlab. As the models have an equal number of parameters, model comparisons were performed by directly comparing coefficient of determination (R^2^) values. R^2^ values were estimated using linear regression.

### Identification of Laminar Borders and Analysis of RF Alignment

To ensure that the array was positioned orthogonal to the cortical surface, we used as criteria the vertical alignment of the mapped RF at each contact (see **Supplementary Fig. 1b**), as well as the similarity in the orientation tuning curves recorded at each contact. The array was removed from cortex and repositioned, if significant RF misalignments across contacts were detected. The degree of RF misalignment was also quantified for each penetration as described in the Supplementary Results (see Analysis of RF Alignment).

The borders between the granular layer (4C) and supra- and infragranular layers were determined by applying current source density (CSD) analysis, using the kernel CSD method^58^, to the band-pass filtered (1-100 Hz) and trial averaged (n=400) continuous voltage traces evoked by a brief full-field luminance increment (100ms, every 400ms, 1-89cd m^−2^; **Supplementary Fig. 1a**). As previously established^59^, the earliest current sink corresponds to the granular layer, and its borders with the supra- and infra-granular layers can be determined from the reversals from current sink to current source above and below the granular layer, respectively.

### Statistical Analysis

Statistical p-values refer to either independent sample or one sample two-tailed T-tests. For the within layer comparisons (**Fig. 2b_4_**), where the expected effect direction was known, one-tailed t-tests are reported. When deviations from normality were detected using QQ-plots (RF size analysis), the T-tests were augmented with Mann-Whitney U-test. The variances of statistically compared groups were not significantly different (Levene’s test P > 0.2 for RF size comparisons; F-test p > 0.17 for response amplitude comparisons). Unless otherwise specified, for all groups, mean±standard error (s.e.m.) of the mean is reported.

### Suppression Index

The Suppression Index (SI) in **Fig. 3a_2_-a_3_** was computed as follows: SI_no-laser_ = (R_C-*no*-*laser*_ - R_CS-*no*-*laser*_)/R_C-*no*-*laser*_. SI_laser_ = (R_C-*no*-*laser*_ - R_CS-*laser*_)/R_C-*no*-*laser*_, where R_C-*no*-*laser*_ is the response to a stimulus confined to the RF (the peak of the summation curve) in the no-laser condition, R_CS-*no*-*laser*_ is the response to the stimulus covering the RF and surround in the no-laser condition (the proximal surround only for the measurements in **Fig. 3a_2_**, and the full extent of the surround for the measurements in **Fig. 3a_3_**), and R_CS-*laser*_ is the response to the stimulus covering the RF and surround in the laser condition.

### Histology

On completion of the recording session, the animal was perfused transcardially with 2-4% paraformaldehyde in 0.1M phosphate buffer. The occipital pole was frozen-sectioned at 40μm, tangentially to the cortical surface (n=2 brains), or sagittally (n=1). GFP label in V2 and V1 and DiI tracks were visualized under fluorescence to ascertain injection sites were confined to V2, and electrode penetrations were targeted to regions expressing GFP (**Fig. 1d,e_2_**). Electrode penetrations from regions with low GFP expression were eliminated from analysis. Sections were counterstained with DAPI (Sigma-Aldrich, St. Louis, MO) to identify V1/V2 border and cortical layers (**Fig. 1d_2_**).

### Network Model

The network mechanisms underlying the observed effects of feedback inactivation were investigated using the model of Schwabe et al^37^. We used exactly the same recurrent network architecture and parameters as in the original published model, which was shown to capture several response properties of surround suppression in V1, including contrast-dependent changes in RF size and surround suppression strength.

For model details we refer the reader to the original publication. Briefly, the network model represents two areas of visual cortex, V1 and an extra-striate area, each area simplified to a single cortical layer. A schematic diagram illustrating the basic network architecture is shown in **Fig. 5a**. Each spatial location in the model is represented by coupled local excitatory (E) and inhibitory (I) cells, which act as the basic functional module of the network that incorporates the effects of local recurrent connections. Interactions between these modules are mediated by horizontal and feedback connections. The spatial profile and conduction velocities of horizontal and feedback connections are constrained by existing anatomical and physiological data, according to which feedback connections are spatially more extensive^34^ and have faster conduction velocities^45^ than horizontal connections. Because we are focusing on size-tuning effects in this study, it seemed sufficient to take a very simple local network model with a single inhibitory neuron type. The stimulations were run with 30% contrast, which is equivalent to translating the contrast response functions of the model neurons along the contrast axis. This modification is justified as V1 neurons exhibit a variety of contrast preferences.

### Data Availability

The data will be made available upon reasonable request to the authors.

## ACKNOWLEDGMENTS

We thank Kesi Sainsbury for technical assistance, and Dr. Valentin Dragoi for useful comments on the manuscript. This work was supported by grants from the National Institute of Health (R01 EY026812, R01 EY019743, BRAIN U01 NS099702), the National Science Foundation (IOS 1355075, EAGER 1649923), the University of Utah Research Foundation, The University of Utah Neuroscience Initiative, to A.A., a grant from Research to Prevent Blindness, Inc. to the Department of Ophthalmology, University of Utah, and a postdoctoral fellowship from the Ella and Georg Ehrnrooth Foundation to L.N. The authors declare no competing financial interests.

## AUTHOR CONTRIBUTIONS

L.N., S.M., M.B. and A.A. designed project and collected electrophysiological data. L.N, S.M. and F.F. performed optical imaging and viral injections. L.N. analyzed optogenetic and electrophysiological data. S.M. analyzed optical imaging data and histological expression of GFP label. S.M. and F.F. generated histological figures. L.N. and S.M. built the optogenetic stimulation system. A.A. supervised all aspects of project. L.N., S.M. and A.A. wrote the paper. All authors discussed the results, commented on and approved the final manuscript.

## SUPPLEMENTARY RESULTS

### Analysis of RF Alignment

For each array penetration, we quantified the alignment of the minimum response fields (mRFs) mapped at each contact across the array. For each contact, we calculated the Euclidean distance of the mRF center, mapped at that contact, from the mean of mRF centers in that penetration. For the purpose of this analysis, the mRF center was defined as the center of mass of the thresholded mRF activity map (responses >1SD above baseline; **Supplementary Fig. 1b**). The median Euclidean distance from the mean mRF center of each penetration was 0.124°±0.029°/0.035° (95% CI lower bound/upper bound; bootstrap).

### Control Experiments in Cortex Not Expressing ArchT

For the main experiment, laser intensities were selected based on a control experiment in one animal (n=2 penetration) in cortex not expressing ArchT. Recordings and analysis were otherwise identical to the main experiment. Unsorted, thresholded multi-units were analyzed for this control.

We found light artifacts at relatively low light intensities (63mW/mm^2^; see **Supplementary Fig. 2a**), which, to our surprise, have been used in previous optogenetic experiments. The laser artifacts were qualitatively different in superficial and deep layers: spike-rates were usually increased in superficial layers, but often decreased in deep layers (**Supplementary Fig. 2a**). For granular and infragranular layers, irradiances at or below 43mW/mm^2^ did not produce statistically significant changes in the cells’ size tuning curves (mean RF size±s.e.m no-laser vs. laser: granular 1.84±0.04° vs. 1.76±0.11° p=0.60, n=5; infragranular: 2.75±0.75° vs. 2.50±1.0°, p=0.91, n=2; **Supplementary Fig. 2a-b**). For some contacts in supragranular layers, instead, the laser-on and control curves differed significantly at 43mW/mm^2^ irradiance. Importantly, however, the effect of light on these cells was always a *decrease* in RF diameter, i.e. an effect opposite to that caused by the laser in ArchT expressing cortex (supragranular: 1.57±0.11° vs. 1.24±0.09° p=0.006, n=14 **Supplementary Fig. 2a-b**). These laser-induced artifacts in supragranular cells disappeared at lower laser intensities. Specifically, when we repeated the control RF size analysis including supragranular units at laser irradiances of <43mW/mm^2^, (specifically 19mW/mm^2^; n=12 units for this analysis as we did not have recordings at 19mW/mm^2^ laser irradiance for all units), and granular and infragranular contacts at laser irradiance of 43mW/mm^2^, we found no statistically significant changes in RF diameter (mean±s.e.m no-laser vs. laser: 2.1±0.20° vs. 1.8±0.18°, p=0.36, n=12, mean decrease 6.68±6.96%; **Supplementary Fig. 2c**) or response amplitude in the proximal surround (no-laser vs. laser: 60.9±12.3 vs. 59.6±13.8 spikes/s, p=0.72, n=12, mean decrease 7.7±8.30%; **Supplementary Fig. 2d**).

The analysis reported in the Results (**Figs. 2–3**) was performed for laser intensities up to 43mW/mm^2^, because the laser artifacts induced in some supragranular cells at this intensity could not account for the observed effects of feedback inactivation (as these artifacts caused decreases rather than increases in RF size). However, to further corroborate that our results of feedback inactivation could not be attributed to laser-induced artifacts, we repeated all the main analyses of data recorded in ArchT-expressing cortex, after excluding supragranular units which showed inactivation effects at laser irradiances >19mW/mm^2^, i.e. irradiance levels that had produced artifacts in some supragranular layer cells in control cortex. The results of this analysis were qualitatively and quantitatively similar to those of the original analysis, as detailed below.

#### Analysis of Data in Cortex Expressing ArchT, Excluding Supragranular Cells Showing Inactivation Effects at >19mW/mm^2^ Irradiance

Mean RF diameter was significantly smaller with intact feedback, compared to when feedback was inactivated (mean±s.e.m no-laser vs. laser: 1.24±0.11° vs. 1.83±0.17°, p=0.007, n=26; **Supplementary Fig. 3a**), with a mean increase of 59.3±13.0% (p<0.001). As for the original analysis (**Fig. 3a**), stimuli extending into the proximal surround evoked larger neuronal responses (no-laser vs. laser: 42.0±15.4 vs. 51.8±21.5 spikes/s, mean increase 30.0±6.34%, p<0.01; **Supplementary Fig. 3b_1_**), and therefore less surround suppression, when feedback was inactivated compared to when feedback was intact. Laser stimulation reduced the suppression index (SI) for stimuli covering the RF and proximal surround (SI no-laser vs. laser: 0.18±0.03 vs. -0.02±0.06, p<0.01; **Supplementary Fig. 3b_2_**). In contrast, the response (no-laser vs. laser: 13.1±2.63 vs. 13.3 ±2.73 spikes/s, mean spike-rate increase 10.6±11.2%, p=0.37) and SI (no-laser vs. laser: 0. 57±0.05 vs. 0.55±0.06; **Supplementary Fig. 3b_3_**) evoked by stimuli extending into the more distal surround were unchanged by feedback inactivation. Stimuli confined to the neurons’ RF evoked lower responses in the laser condition (41.1±19.2 spikes/s) vs. the non-laser condition (50.1±17.6 spikes/s, mean reduction 30.3±6.34%, p<0.001; **Supplementary Fig. 3c**). We conclude that increased RF diameter and reduced surround suppression indeed resulted from inactivating V2 feedback to V1, and were not caused by laser-induced heat artifacts.

None of the units recorded in the control experiment showed reduced response at the irradiances used for the analysis of data in **Fig. 4**. Thus, we are confident that the general response suppression for small and large stimuli observed in the data reported in **Fig. 4** resulted from inactivating feedback axons.

### Analysis of RF Size Increase Induced by Noise

We performed an analysis to exclude the possibility that the increased RF size after feedback inactivation could arise from noise in the spatial summation data. For each cell, we first generated a size-tuning curve from the fitted ROG or DOG model, depending on which model better fitted the cell’s response. For each cell, we then generated a “noisy” curve by independently sampling for each presented stimulus diameter 10 responses from a Poisson distribution having the same mean as the “real” fitted curve at those diameters. These 10 sampled responses per stimulus diameter were then averaged to produce a noisy version of the real curve (**Supplementary Figure 4a**), and the peak of the noisy curve was taken as the RF size. This procedure was repeated 10,000 times for each cell. For each repetition, we computed the percent increase in RF size between the real curve and the noisy curve for the cell, as done for the recorded data. Finally, we produced a distribution of the average percent increase in RF size across the population of cells in this simulation. The median increase in RF size in the bootstrapped distribution was 3.6%, as opposed to the 56% increase seen in the real data (**Supplementary Figure 4b**). We thus conclude that feedback inactivation increases RF size significantly more than would be expected from noise.

### Analysis of RF Surround Size

For the population of cells showing an increase in RF diameter when feedback was inactivated, we found no changes in the size of the surround field (see Methods for definition). Average surround diameter in the no-laser vs. laser condition was 4.71±0.43° vs. 5.38±2.77° (p=0.33).

### Control Analysis for Laser Stimulation Time

Inactivation of axon terminals using ArchT can, counter intuitively, facilitate synaptic transmission for prolonged light pulses, while ArchT is consistently suppressive for pulse widths of ≤ 200ms. Thus, we repeated our analysis by focusing only on the first 200ms of the response. We found no qualitative differences between the original analysis and the short time-scale analysis. Consistent with the original analysis, RF diameter was increased when feedback was inactivated (no-laser vs. laser: 1.14±0.07° vs. 1.67±0.24, p<0.05, n=19 units producing reliable responses within the initial 200ms), responses to stimuli confined to the RF were significantly reduced (no-laser vs. laser: 26.1±8.89 vs. 21.6±10.3 spikes/s, mean spike-rate reduction 45.1±8.62%, p<0.001), and responses to stimuli covering the RF and proximal surround were increased (mean spike-rate increase 67.6±34.0%, p<0.06). We conclude that the observed laser-induced effects reflect suppressed, rather than facilitated, V2 feedback activity.

### Phenomenological modeling

To gain insights into the mechanisms underlying feedback inactivation effects, we fitted ROG and DOG models to the data (see Methods) and determined which model parameters were mostly affected by feedback inactivation. To this goal, we selected for each cell the model that provided the best fit to its size tuning measurements (based on the averaged R^2^ over laser and no-laser conditions) in the no-laser condition, and then allowed one parameter at a time to vary with feedback inactivation, while the remainder of the model parameters were held fixed to their original values for the no-laser condition. The model in which feedback inactivation modified the spatial extent of the center excitation (w_c_ in equations 1–3) provided the best fit for 45% of the cells (top left panel in **Supplementary Fig. 5a**). This model could capture the increase in RF size and response reduction for small stimuli, seen in the data after feedback inactivation, but failed to capture increases in response for stimuli extending into the surround. In contrast, a model in which feedback inactivation modified the spatial extent of the surround inhibition (w_s_ in equations 1,2,4) could capture changes in response for stimuli extending into the surround, but failed to capture changes in RF size (top right panel in **Supplementary Fig. 5a**). Moreover, none of the single parameter models could capture the co-occurrence of response reduction to small stimuli and response increase in the proximal surround often seen in the data (**Supplementary Fig. 5a**). We next allowed two parameters at a time to vary with feedback inactivation, while holding the rest constant, and performed this analysis for all possible combinations of parameter pairs. A model in which the spatial extent and gain of the center excitatory mechanism (w_c_ and g_c_, respectively, in equations 1–3) were simultaneously varied provided the best fit for the largest proportion (30%) of cells (**Supplementary Fig. 5b**). A model in which the spatial extent of both the excitatory (w_c_) and inhibitory (w_s_ in equations 1,2,4) mechanisms were varied performed second best, providing the best fit for 21% of cells. For the model in which wc and gc were free parameters, the wc parameter, describing the spatial extent of the center excitation, typically increased as a result of feedback inactivation, paralleling the RF size increase seen in the data (**Supplementary Fig. 5c**). This model, was also able to account for a variety changes in response amplitude, as seen in the data, including cooccurrence of reduced response for stimuli inside the RF and increased response for stimuli in the proximal surround (*solid green curve* in the inset of **Supplementary Fig. 5b**), as well as overall reduced response throughout the entire spatial summation curve (*dashed green curve* in inset of **Supplementary Fig. 5b**). The model in which wc and ws were free parameters could also account for co-occurrence of reduced response for stimuli inside the RF and increased response for stimuli in the proximal surround. Previously, a model in which the gain of the center and surround mechanisms (g_c_ and g_s_, respectively) were free parameters was found to provide a good description of contrast-dependent changes in RF size^1^, which resemble the effects of feedback inactivation we have observed in some cells, especially at high laser intensity. Such a model provided the best fit for only 2 cells in our population (**Supplementary Fig. 5b**), as it failed to account, at least in a straightforward way, for the co-occurrence of response reduction to stimuli in the RF and response increase in the proximal surround, often seen in the data.

However, we found that the different models performed similarly when compared based on the coefficient of determination (R^2^) distributions, rather than fraction of cells best fit by each model (**Supplementary Fig. 5d**).

---

**Supplementary Figure 1.**
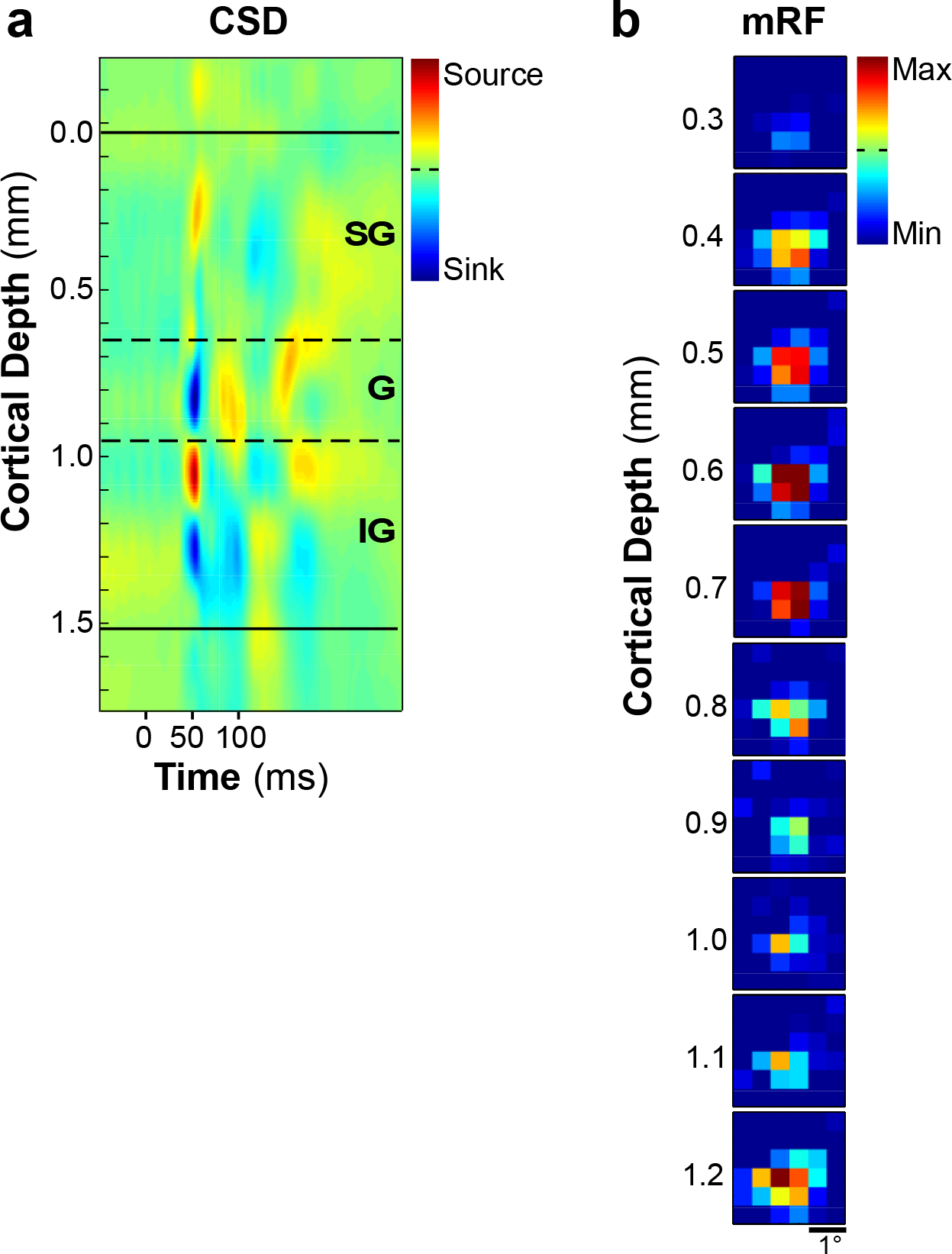
Recordings of CSD and minimum RF (mRF) ensure linear array spans all cortical layers, and is positioned normal to the cortical surface. **(a)** Current source density (CSD) analysis of local field potential (LFP), used to determine cortical layers and ensure contacts span the full extent of the cortical sheet. **(b)** mRF mapping (see Methods) across contacts through the depth of a single array penetration in V1. Hot spots (regions of max spiking rate) are aligned across contacts, confirming the array is positioned normal to the V1 surface. We also quantified the mRF alignment across all penetrations (see Supplementary Results, *Analysis of RF alignment*). *SG*: Supragranular layers, *G*: Granular layer, *IG*: Infragranular layers.

**Supplementary Figure 2.**
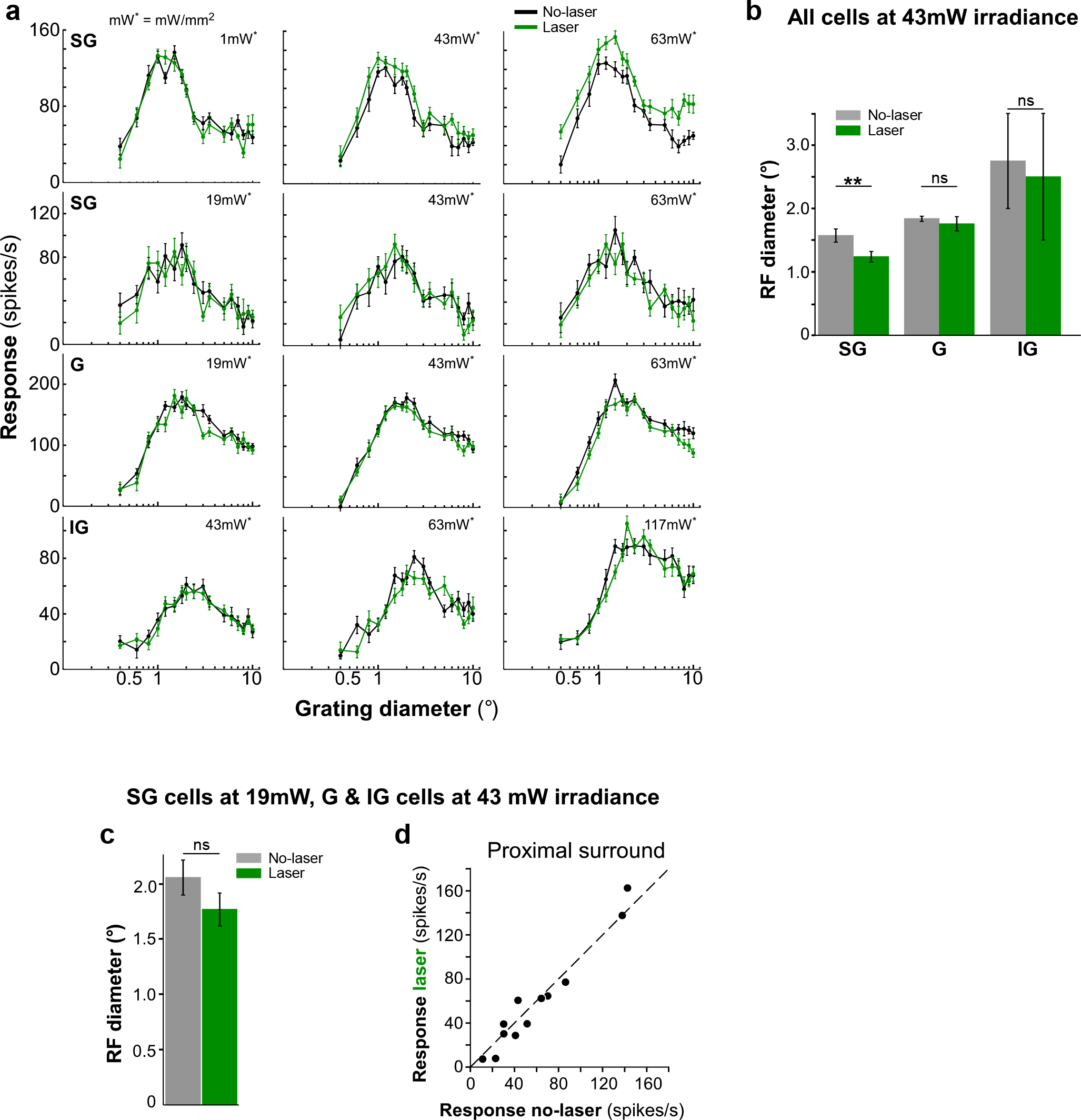
Analysis in control cortex not expressing ArchT. **(a)** Spatial summation curves for two supragranular (SG), a granular (G), and an infragranular (IG) example cells with (*green*) and without (*black*) laser stimulation at different light intensities (irradiance indicated). **(b)** Mean RF diameter, with and without laser stimulation, for the whole population of multiunits at laser irradiance of 43mW/mm^2^, grouped by layer location. **(c)** Mean RF diameter, with and without laser stimulation, for the whole population of cells including SG cells treated at laser irradiance of 19mW/mm^2^ and G and IG cells at laser irradiance of 43mW/mm^2^ **(d)** Response with and without laser for stimuli involving the RF and proximal surround, for the same population of cells as in (c).

**Supplementary Figure 3.**
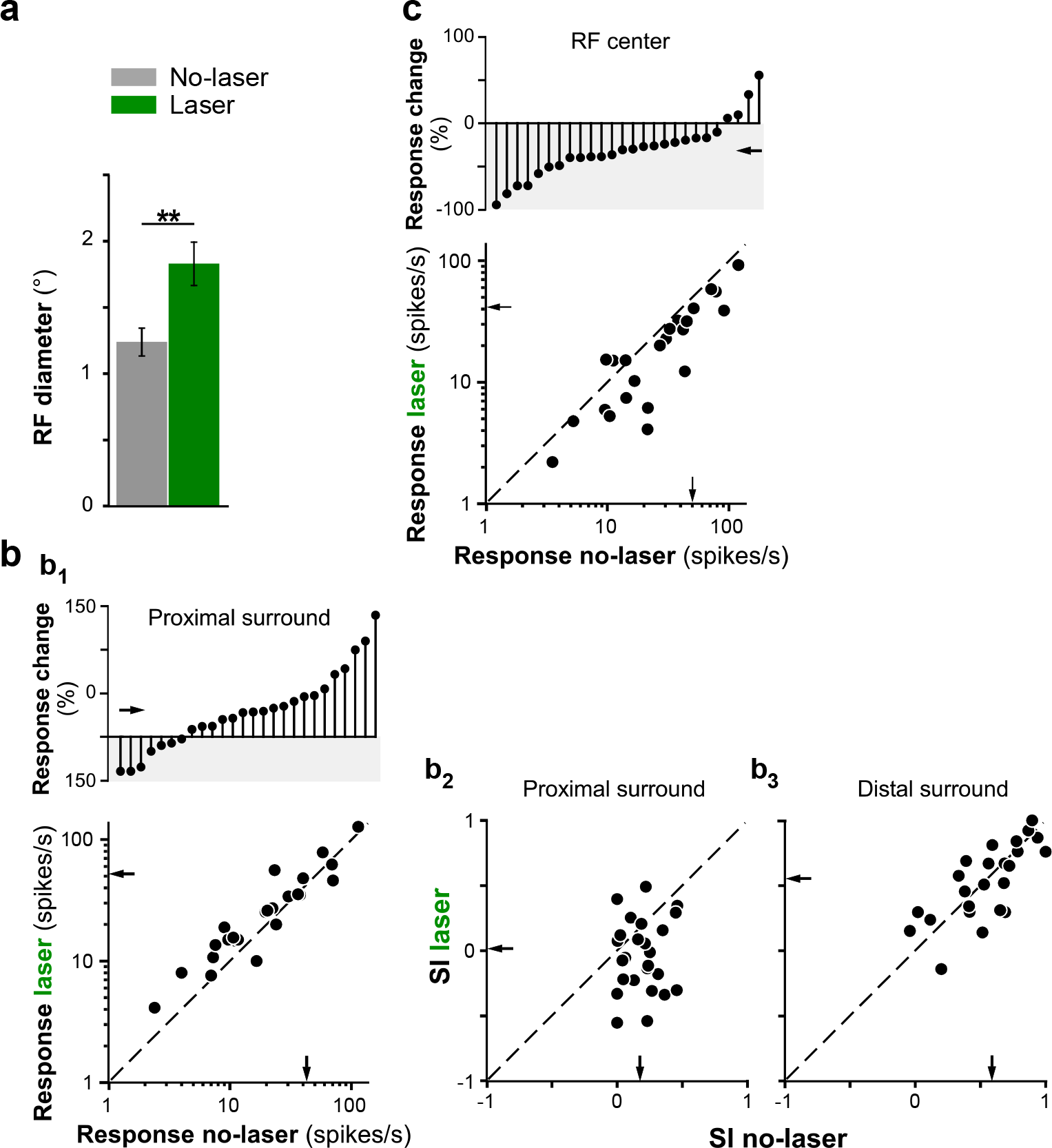
Analysis in cortex expressing ArchT, excluding supragranular cells showing inactivation effects at laser irradiance >19 mW/mm^2^. **(a)** Mean RF diameter with and without laser stimulation. **(b)** Changes in surround suppression with V2 feedback inactivated. **(b_1_)** BOTTOM: response with and without laser for stimuli involving the RF and proximal surround. TOP: Cell-by-cell percent response change induced by laser stimulation, for stimuli extending into the proximal surround. Upward stems: increase in response with laser stimulation; downward stems (*gray shading)*: decrease in response with laser stimulation. **(b_2_)** SI with and without laser for stimuli extending into the proximal surround. **(b_3_)** Same as (b_2_) but for stimuli extending into the distal surround. **(c)** BOTTOM: response with and without laser for stimuli confined to the RF. TOP: Cell-by-cell percent response change induced by laser stimulation, for stimuli confined to the RF. *Arrows in (b-c)*: means.

**Supplementary Figure 4.**
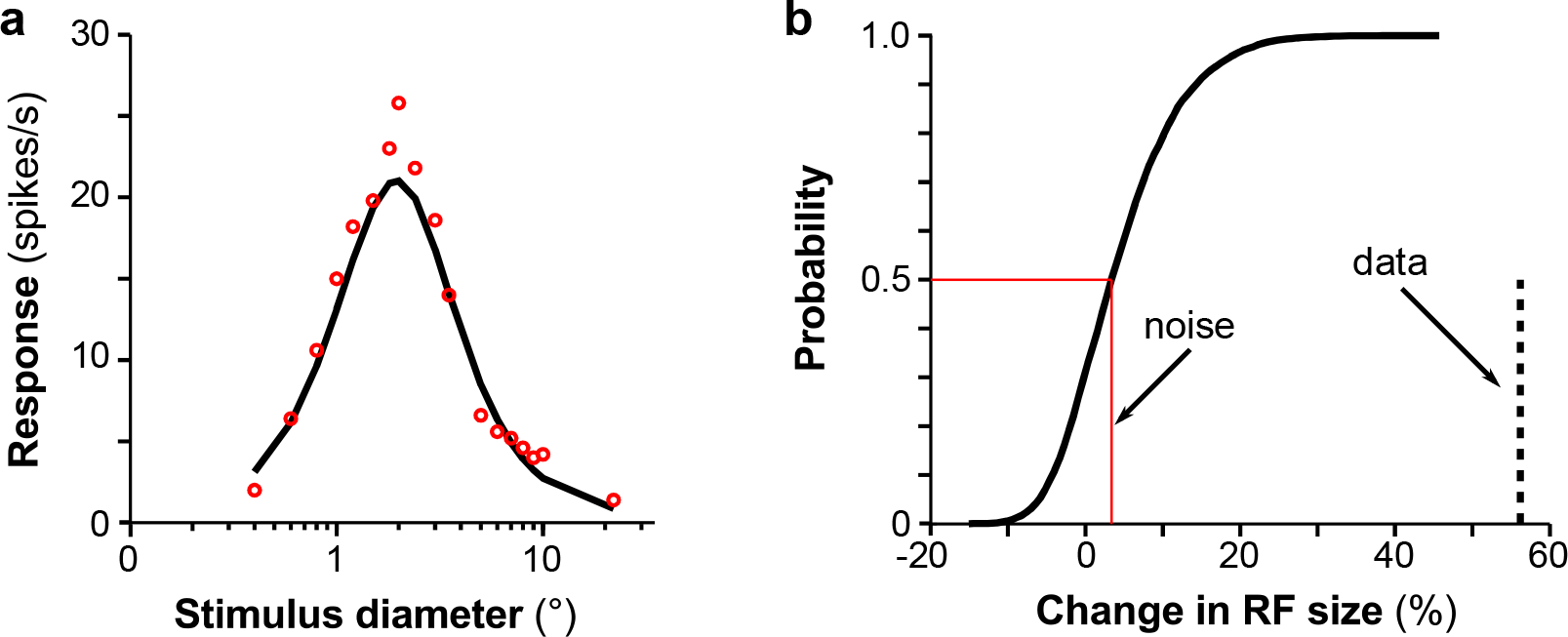
Analysis of RF size increase induced by noise. **(a)** The *black curve* represents the “real” size tuning curve derived from phenomenological model fits to the spatial summation data of an example V1 cell. The *red circles* represent the simulated response at each stimulus diameter averaged over 10 trials. The response in each trial was obtained by randomly sampling from a Poisson distribution with the same mean as in the “real” curve. **(b)** Cumulative distribution of percent RF size changes expected under the null hypothesis that the real size tuning curve measured with and without laser stimulation were identical, and all changes in RF size were due to noise. RF size change due to noise was computed separately for each cell, based on a “noisy” tuning curve formed by averaging simulated responses in 10 trials, then averaging over the population of 33 cells and repeating the procedure 10,000 times (see Supplementary Results for details).

**Supplementary Figure 5.**
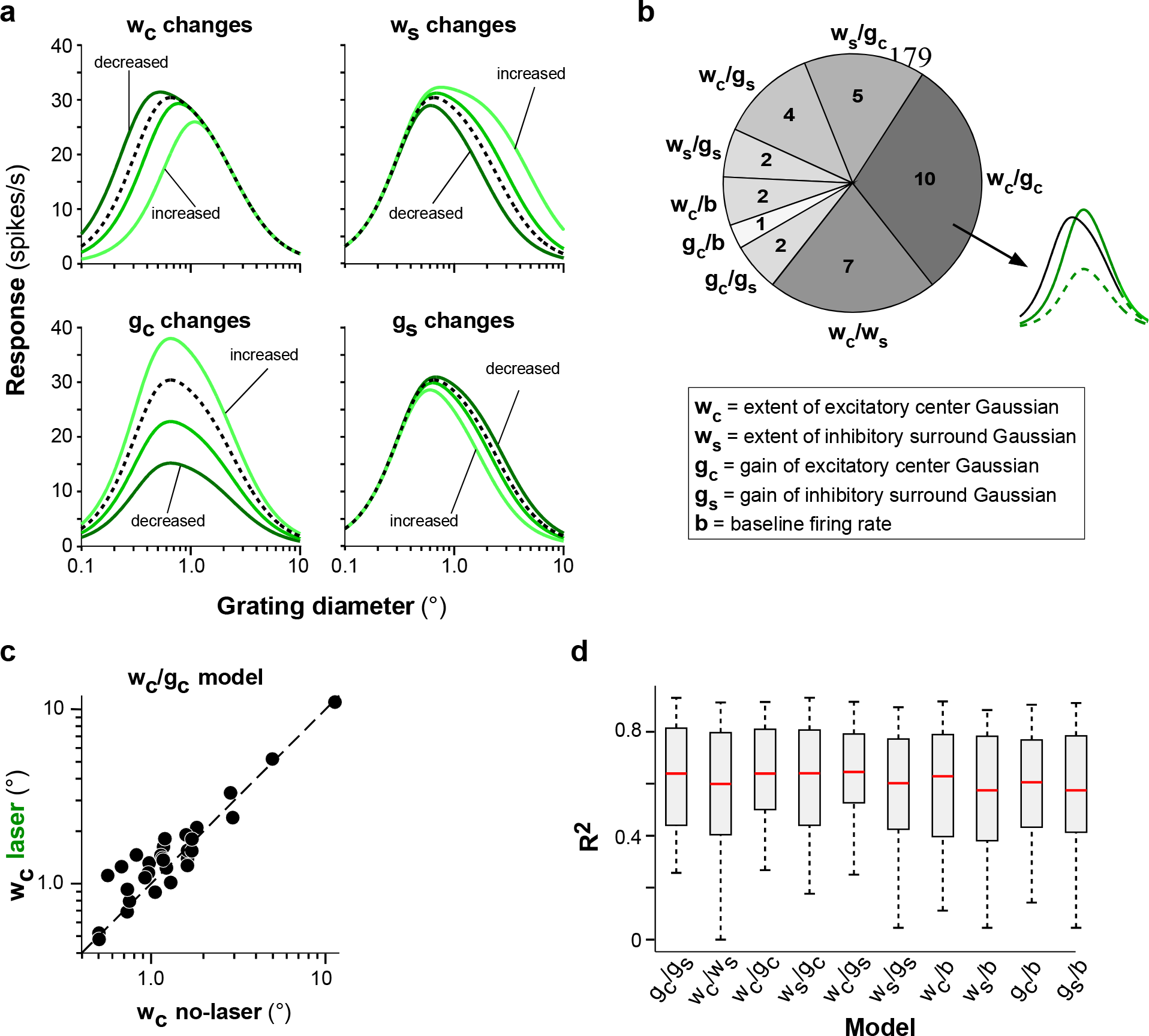
Phenomenological models. **(a)** Effects of feedback inactivation for each of 4 different versions of the phenomenological models (DOG and ROG), in which only one parameter at a time was allowed to vary while keeping the other parameters fixed to their value in the model that provided the best fit to size tuning measurements in the no-laser condition (*black curve*). In each panel, the *black curve* represents the best fit to the spatial summation data in the no-laser condition of the same representative V1 cell; the *green curves* represent changes in the summation curve when only one parameter (the one indicated at the top of the panel) was allowed to change while keeping the others fixed at their values in the black curve. Different shadings of green indicate different levels of feedback activity, achieved in the model by scaling the free parameter by 0.75 (*darkest green*), 1.25 or 2 (*lightest green*), in all panels, but the top left, for which scaling factors of w_c_ were 0.5, 0.75 and 1.25, respectively. None of the models could account for the co-occurrence of RF size increase and increased response in the proximal surround, as often seen in the data (see **Fig. 2a**). **(b)** Proportion of cells for wich each of the two-parameter models (in which only two parameters at a time, i.e. those indicated next to each wedge, were allowed to vary) provided the best fit to the inactivation data. *Numbers* in each wedge indicate the number of cells best fitted by each model. The *inset* indicates the two main effects of feedback inactivation on size tuning that the w /g model could account for. **(c)** Scatterplot of the spatial extent of the center Gaussian mechanism (w_c_) in the no-laser vs. the laser condition, in the w_c_/g_c_ model. Feedback inactivation in this model generally increased the spatial extent of the center Gaussian. **(d)** Distribution of R^2^ for each of the two-parameter models. *Red lines*: medians.

## REFERENCES

1. Van Essen, D. C. & Maunsell, J. H. R. Hierarchical organization and functional streams in the visual cortex. Trends Neurosci. 6, 370–375 (1983).

2. McAdams, C. J. & Reid, C. R. Attention modulates the responses of simple cells in monkey primary visual cortex. J. Neurosci. 25, 11023–11033 (2005).

3. Luck, S. J., Chelazzi, L., Hillyard, S. A. & Desimone, R. Neural mechanims of spatial selective attention in areas V1, V2 and V4 of macaque visual cortex. J. Neurophysiol. 77, 24–42 (1997).

4. Rao, R. P. & Ballard, D. H. Predictive coding in the visual cortex: a functional interpretation of some extra-classical receptive-field effects. Nat. Neurosci. 2, 79–87 (1999).

5. Angelucci, A., Bijanzadeh, M., Nurminen, L., Federer, F., Merlin, S. & Bressloff, P. C. Circuits and mechanisms for surround modulation in visual cortex. Ann. Rev. Neurosci. 40, In Press (2017).

6. Angelucci, A. & Bressloff, P. C. The contribution of feedforward, lateral and feedback connections to the classical receptive field center and extra-classical receptive field surround of primate V1 neurons. Prog. Brain Res. 154, 93–121 (2006).

7. McAdams, C. J. & Maunsell, J. H. Effects of attention on orientation-tuning functions of single neurons in macaque cortical area V4. J. Neurosci. 19, 431–441 (1999).

8. Sundberg, K. A., Mitchell, J. F. & Reynolds, J. H. Spatial attention modulates center-surround interactions in macaque visual area V4. Neuron 61, 952–963 (2009).

9. Roberts, M. J., Delicato, L. S., Herrero, J., Gieselmann, M. A. & Thiele, A. Attention alters spatial integration in macaque V1 in an eccentricity dependent manner. Nat. Neurosci. 10, 1483–1491 (2007).

10. Shushruth, S., Ichida, J. M., Levitt, J. B. & Angelucci, A. Comparison of spatial summation properties of neurons in macaque V1 and V2. J. Neurophysiol. 102, 2069–2083 (2009).

11. Allman, J., Miezin, F. & Mc Guinness, E. Stimulus specific responses from beyond the classical receptive field: Neurophysiological mechanisms for local-global comparisons in visual neurons. Ann. Rev. Neurosci. 8, 407–430 (1985).

12. Nurminen, L., Kilpelainen, M., Laurinen, P. & Vanni, S. Area summation in human visual system: psychophysics, fMRI, and modeling. J. Neurophysiol. 102, 2900–2909 (2009).

13. Blakemore, C. & Tobin, E. A. Lateral inhibition between orientation detectors in the cat’s visual cortex. Exp. Brain Res. 15, 439–440 (1972).

14. Cavanaugh, J. R., Bair, W. & Movshon, J. A. Nature and interaction of signals from the receptive field center and surround in macaque V1 neurons. J. Neurophysiol. 88, 2530–2546 (2002).

15. Sceniak, M. P., Hawken, M. J. & Shapley, R. M. Visual spatial characterization of macaque V1 neurons. J. Neurophysiol. 85, 1873–1887 (2001).

16. Hubel, D. H. & Wiesel, T. N. Receptive fields and functional architecture in two nonstriate visual areas (18 and 19) of the cat. J. Neurophysiol. 28, 229–289 (1965).

17. Angelucci, A. & Shushruth, S. Beyond the classical receptive field: surround modulation in primary visual cortex. In: The new visual neurosciences (ed^(eds Chalupa, L. M., Werner, J. S.). MIT press (2013).

18. Van den Bergh, G., Zhang, B., Arckens, L. & Chino, Y. M. Receptive-field properties of V1 and V2 neurons in mice and macaque monkeys. J. Comp. Neurol. 518, 2051–2070 (2010).

19. Olshausen, B. A. & Field, D. J. Emergence of simple-cell receptive field properties by learning a sparse code for natural images. Nature 381, 607–609 (1996).

20. Vinje, W. E. & Gallant, J. L. Natural stimulation of the nonclassical receptive field increase information transmission efficiency in V1. J. Neurosci. 22, 2904–2915 (2002).

21. Nurminen, L. & Angelucci, A. Multiple components of surround modulation in primary visual cortex: multiple neural circuits with multiple functions? Vision research 104, 4756 (2014).

22. Schwartz, O. & Simoncelli, E. P. Natural signal statistics and sensory gain control. Nat. Neurosci. 4, 819–825 (2001).

23. Hupé, J. M., James, A. C., Payne, B. R., Lomber, S. G., Girard, P. & Bullier, J. Cortical feedback improves discrimination between figure and background by V1, V2 and V3 neurons. Nature 394, 784–787 (1998).

24. Bardy, C., Huang, J. Y., Wang, C., Fitzgibbon, T. & Dreher, B. “Top-down” influences of ispilateral or contralateral postero-temporal visual cortices on the extra-classical receptive fields of neurons in cat’s striate cortex. Neurosci. 158, 951–968 (2009).

25. Nassi, J. J., Lomber, S. G. & Born, R. T. Corticocortical feedback contributes to surround suppression in V1 of the alert primate. J. Neurosci. 33, 8504–8517 (2013).

26. Zhang, S., et al. Long-range and local circuits for top-down modulation of visual cortex processing. Science 345, 660–665 (2014).

27. Hupé, J. M., James, A. C., Girard, P. & Bullier, J. Response modulations by static texture surround in area V1 of the macaque monkey do not depend on feedback connections from V2. J. Neurophysiol. 85, 146–163 (2001).

28. Wang, C., Huang, J. Y., Bardy, C., FitzGibbon, T. & Dreher, B. Influence of ‘feedback’ signals on spatial integration in receptive fields of cat area 17 neurons. Brain Res. 1328, 34–48 (2010).

29. Sandell, J. H. & Schiller, P. H. Effect of cooling area 18 on striate cortex cells in the squirrel monkey. J. Neurophysiol. 48, 38–48 (1982).

30. Han, X., et al. A high-light sensitivity optical neural silencer: development and application to optogenetic control of non-human primate cortex. Front. Syst. Neurosci. 5, 18 (2011).

31. Stujenske, J. M., Spellman, T. & Gordon, J. A. Modeling the Spatiotemporal Dynamics of Light and Heat Propagation for In Vivo Optogenetics. Cell Rep. 12, 525–534 (2015).

32. Rockland, K. S. The organization of feedback connections from area V2 (18) to V1 (17). In: Primary Visual Cortex in Primates (ed^(eds Peters, A., Rockland, K. S.). Plenum Press (1994).

33. Federer, F., Merlin, S. & Angelucci, A. Anatomical and functional specificity of V2-to-V1 feedback circuits in the primate visual cortex. Soc. Neurosci. Abstr. Online., 699.602 (2015).

34. Angelucci, A., Levitt, J. B., Walton, E., Hupé, J. M., Bullier, J. & Lund, J. S. Circuits for local and global signal integration in primary visual cortex. J. Neurosci. 22, 8633–8646. (2002).

35. Mahn, M., Prigge, M., Ron, S., Levy, R. & Yizhar, O. Biophysical constraints of optogenetic inhibition at presynaptic terminals. Nat Neurosci 19, 554–556 (2016).

36. Sceniak, M. P., Ringach, D. L., Hawken, M. J. & Shapley, R. Contrast’s effect on spatial summation by macaque V1 neurons. Nat. Neurosci. 2, 733--739 (1999).

37. Schwabe, L., Obermayer, K., Angelucci, A. & Bressloff, P. C. The role of feedback in shaping the extra-classical receptive field of cortical neurons: a recurrent network model. J. Neurosci. 26, 9117–9129 (2006).

38. Adesnik, H., Bruns, W., Taniguchi, H., Huang, Z. J. & Scanziani, M. A neural circuit for spatial summation in visual cortex. Nature 490, 226–231 (2012).

39. Henry, C. A., Joshi, S., Xing, D., Shapley, R. M. & Hawken, M. J. Functional characterization of the extraclassical receptive field in macaque V1: contrast, orientation, and temporal dynamics. J. Neurosci. 33, 6230–6242 (2013).

40. Webb, B. S., Dhruv, N. T., Solomon, S. G., Taliby, C. & Lennie, P. Early and late mechanisms of surround suppression in striate cortex of macaque. J. Neurosci. 25, 11666–11675 (2005).

41. Bijanzadeh, M., Nurminen, L., Merlin, S. & Angelucci, A. Distinct laminar processing of local and global context in primate primary visual cortex. BioRxiv, https://doi.org/10.1101/171793 (2017).

42. Alitto, H. J. & Usrey, W. M. Origin and dynamics of extraclassical suppression in the lateral geniculate nucleus of the macaque monkey. Neuron 57, 135–146 (2008).

43. Sceniak, M. P., Chatterjee, S. & Callaway, E. M. Visual spatial summation in macaque geniculocortical afferents. J. Neurophysiol. 96, 3474–3484 (2006).

44. Sato, T. K., Hausser, M. & Carandini, M. Distal connectivity causes summation and division across mouse visual cortex. Nat. Neurosci. 17, 30–32 (2014).

45. Girard, P., Hupé, J. M. & Bullier, J. Feedforward and feedback connections between areas V1 and V2 of the monkey have similar rapid conduction velocities. J. Neurophysiol. 85, 1328–1331. (2001).

46. Bair, W., Cavanaugh, J. R. & Movshon, J. A. Time Course and Time-Distance Relationships for Surround Suppression in Macaque V1 Neurons. J. Neurosci. 23, 7690–7701 (2003).

47. Nassi, J. J., Gomez-Laberge, C., Kreiman, G. & Born, R. T. Corticocortical feedback increases the spatial extent of normalization. Front. Syst. Neurosci. 8, 105 (2014).

48. Rubin, D. B., Van Hooser, S. D. & Miller, K. D. The stabilized supralinear network: a unifying circuit motif underlying multi-input integration in sensory cortex. Neuron 85, 402–417 (2015).

49. Ito, M. & Gilbert, C. D. Attention modulates contextual influences in the primary visual cortex of alert monkeys. Neuron 22, 593–604 (1999).

50. Federer, F., Ichida, J. M., Jeffs, J., Schiessl, I., McLoughlin, N. & Angelucci, A. Four projections streams from primate V1 to the cytochrome oxidase stripes of V2. J. Neurosci. 29, 15455–15471 (2009).

51. Schiessl, I. & McLoughlin, N. Optical imaging of the retinotopic organization of V1 in the common marmoset. Neuroimage 20, 1857–1864 (2003).

52. Rossant, C., et al. Spike sorting for large, dense electrode arrays. Nat. Neurosci. 19, 634–641 (2016).

53. Kadir, S. N., Goodman, D. F. & Harris, K. D. High-dimensional cluster analysis with the masked EM algorithm. Neural. Comput. 26, 2379–2394 (2014).

54. Ringach, D. L., Shapley, R. M. & Hawken, M. J. Orientation selectivity in macaque V1: diversity and laminar dependence. J. Neurosci. 22, 5639–5651 (2002).

55. Tailby, C., Solomon, S. G., Peirce, J. W. & Metha, A. B. Two expressions of “surround suppression” in V1 that arise independent of cortical mechanisms of suppression. Visual neuroscience 24, 99–109 (2007).

56. Hunter, J. D. Matplotlib: A 2D graphics environment. Comput. Sci. Eng, 9, 90–95 (2007).

57. van der Walt, S., Colbert, S. C. & Varoquaux, G. The NumPy Array: A Structure for Efficient Numerical Computation. Comput. Sci. Eng. 13, 22–30 (2011).

58. Potworowski, J., Jakuczun, W., Leski, S. & Wojcik, D. Kernel current source density method. Neural. Comput. 24, 541–575 (2012).

59. Schroeder, C. E., Mehta, A. D. & Givre, S. J. A spatiotemporal profile of visual system activation revealed by current source density analysis in the awake macaque. Cereb. Cortex 8, 575–592 (1998).

## REFERENCES

1. Cavanaugh, J. R., Bair, W. & Movshon, J. A. Nature and interaction of signals from the receptive field center and surround in macaque V1 neurons. J. Neurophysiol. 88, 2530–2546 (2002).

